# Revisiting the effect of pharmaceuticals on transmission stage formation in the malaria parasite *Plasmodium falciparum*

**DOI:** 10.1101/2021.11.18.469097

**Authors:** Basil T. Thommen, Armin Passecker, Tamara Buser, Eva Hitz, Till S. Voss, Nicolas M. B. Brancucci

**Author notes:** **Correspondence:** Till Voss, Nicolas Brancucci. These authors have contributed equally to this work and share last authorship (Till S. Voss, Nicolas M. B. Brancucci).

## Abstract

Malaria parasites rely on specialized stages, called gametocytes, to ensure human-to-human transmission. The formation of these sexual precursor cells is initiated by commitment of blood stage parasites to the sexual differentiation pathway. *Plasmodium falciparum*, the most virulent of six parasite species infecting humans, employs nutrient sensing to control the rate at which sexual commitment is initiated, and the presence of stress-inducing factors, including antimalarial drugs, has been linked to increased gametocyte production *in vitro* and *in vivo*. These observations suggest that therapeutic interventions may promote gametocytogenesis and malaria transmission. Here, we engineered a *P. falciparum* reporter line to quantify sexual commitment rates after exposure to antimalarials and other pharmaceuticals commonly prescribed in malaria-endemic regions. Our data reveal that some of the tested drugs indeed have the capacity to elevate sexual commitment rates *in vitro*. Importantly, however, these effects are only observed at drug concentrations that inhibit parasite survival and only rarely result in a net increase of gametocyte production. Using a drug-resistant parasite reporter line, we further show that the gametocytogenesis-promoting effect of drugs is linked to general stress responses rather than to compound-specific activities. Altogether, we provide conclusive evidence for the absence of mechanistic links between the regulation of sexual commitment and the activity of commonly used pharmaceuticals *in vitro*. Our data hence contradict scenarios in which therapeutic interventions would promote the spread of drug-resistant parasites or malaria transmission in general.

## Introduction

Causing an estimated 200 million clinical cases and more than 400′000 deaths annually, malaria represents one of the major threats to global public health (1). Malaria parasites resistant to current drug interventions, including the frontline artemisinin-based combination therapies (ACTs), are emerging and endanger malaria eradication campaigns (2). Among the six *Plasmodium* species infecting humans, *P. falciparum* is the most virulent and accounts for the majority of severe and lethal malaria cases (1). Following the injection of sporozoites by an infected *Anopheles* mosquito, parasites reproduce within hepatocytes before initiating the symptomatic phase of infection in the human blood. The latter is characterized by continuous rounds of erythrocyte invasion, asexual replication, host cell rupture, and the release of merozoites ready to invade new red blood cells (RBCs). During each of these 48-hour long intra-erythrocytic replication cycles, a small subset of parasites switches away from asexual replication and instead commits to sexual development, resulting in the formation of transmissible gametocytes (3). *P. falciparum* gametocytes sequester in deep tissue, including the bone marrow parenchyma, where they undergo a series of developmental steps (I-V) before re-entering the blood stream as mature and transmission-competent stage V gametocytes after 10-12 days (4, 5). While gametocytes represent the only cell type that is infectious to mosquitoes, they are non-replicative. Investments into gametocytogenesis thus come at the expense of reduced vegetative growth and *P. falciparum* employs sophisticated mechanisms to regulate this trade-off between within-host replication and between-host transmission (6).

Sexual commitment requires expression of AP2-G – a member of the ApiAP2 family of DNA-binding factors (7, 8). This master regulator of sexual commitment primes asexually replicating parasites to produce sexually committed ring stage progeny that exit the cell cycle and undergo gametocyte development (3). In addition to this mechanism referred to as ‘next cycle conversion’, the immediate induction of gametocytogenesis in ring stages has also been observed, albeit this ‘same cycle conversion’ is induced at a low rate and has so far only been described *in vitro* (9). In the predominant NCC route, the decision of whether to stay within the asexual pathway or commit to the production of gametocytes is made in early schizonts at 36 +/− 4 hours post-erythrocyte invasion (hpi) (10). While AP2-G expression is initiated almost simultaneously, gametocyte differentiation will only start after completion of the current intra-erythrocytic developmental cycle (IDC) and the invasion of sexually committed merozoites into new RBCs (5). On the molecular level, the process of sexual commitment is under epigenetic control. It involves the activity of a number of well-characterized factors, including heterochromatin protein 1 (HP1), histone deacetylase 2 (HDA2) and gametocyte development protein 1 (GDV1) (3). These factors act in concert to control transcriptional activity at the *ap2-g* locus. The *ap2-g* locus is generally kept in a silenced state marked by the presence of histone 3 tri-methylated at lysine 9 (H3K9me3) and the histone code writer and reader proteins HDA2 and HP1, respectively (11–15). In contrast, GDV1 counteracts the silencing of *ap2-g* by evicting HP1 from the *ap2-g* locus, thus triggering AP2-G expression and sexual commitment in schizonts (16–18). Through an auto-regulatory positive feedback loop, AP2-G expression increases further and peaks in the sexually committed ring stage parasites where it binds to and regulates the expression of genes linked to gametocytogenesis (9, 19, 20). Together, these studies revealed that sophisticated epigenetic mechanisms are in place to balance asexual reproduction versus investments into transmission stage formation (3, 6).

*P. falciparum* parasites invest surprisingly little into transmission. In fact, sexual commitment rates (SCRs) in parasite populations typically remain at a low single-digit percentage *in vitro* and *in vivo* (6, 18, 21–23). This reproductive restraint can be lifted under certain circumstances, resulting in greatly enhanced sexual commitment of parasites and conversion rates above 30% under specific *in vitro* conditions (24). These include, but are not restricted to, high parasite densities (25, 26), exposure to *P. falciparum*-conditioned (nutrient-depleted) medium (23, 27–29), endoplasmic reticulum stress (30) or the uptake of extracellular vesicles derived from infected RBCs (iRBCs) (31, 32). These observations contributed to the appreciation that intra-erythrocytic parasites are able to interact with and respond to their environment. Indeed, recent studies revealed that blood stage parasites modulate specific transcriptional programs in response to nutrient availability and other environmental cues (33). Importantly, *P. falciparum* was found to metabolize the host-derived serum lipid lysophosphatidylcholine (lysoPC) in the Kennedy pathway and to induce sexual commitment under conditions that limit activity of this metabolic route (10). Expression of GDV1, the earliest known marker of sexual commitment, is induced at low lysoPC concentrations, implying a direct link between the epigenetic mechanisms that control gametocyte production and parasite nutrient-sensing (24).

In addition, several lines of evidence suggest that some antimalarial drugs may interfere with the process of sexual commitment (34–39). Increased gametocyte production was observed *in vitro* upon treatment with artemisinin, mefloquine, chloroquine, primaquine, atovaquone and piperaquine (37). Furthermore, treatment with sub-curative doses of the widely used drugs chloroquine (36) and sulfadoxine/pyrimethamine (“Fansidar”) (39) have been associated with increased gametocyte production *in vivo*. Using a transgenic reporter cell line for the quantification of sexual conversion rates (40), the Cortés laboratory recently confirmed a gametocytogenesis-inducing effect for the frontline antimalarial dihydroartemisinin (DHA) (38). In this study, short pulses of sub-curative DHA concentrations applied to trophozoites induced sexual commitment *in vitro*, but this effect was not observed when ring or schizont stage parasites were exposed to the drug. Together, these studies raise legitimate concerns about whether therapeutic interventions may promote gametocytogenesis and hence malaria transmission. However, the mechanisms underlying such potential drug-induced increases in gametocyte production are hitherto unknown. One the one hand, antimalarials may promote sexual commitment by inducing general cellular stress responses. On the other hand, it is conceivable that gametocytogenesis may be induced by drug-specific modulation of the molecular process underlying sexual commitment – i.e. independent of the toxic effect of the drug. In this latter scenario, drug-resistant parasites would be expected to shift their investment towards gametocyte formation under drug pressure, which may eventually promote transmission and spread of drug resistance (41). Indeed, strains carrying specific drug resistance mutations have been associated with increased mosquito transmission following antimalarial treatment (42). Furthermore, chloroquine-resistant parasites showed increased gametocytemia and mosquito infectivity following drug treatment compared to chloroquine-sensitive strains (43). Other efforts, however, failed to confirm the specific induction of gametocyte production in drug-resistant parasites after exposure to sub-curative chloroquine and pyrimethamine concentrations *in vitro* (44). Hence, while drug-resistant parasite strains may have an increased transmission potential, it remains unclear whether this would be linked to higher parasite survival rates under drug pressure or truly increased rates of commitment to gametocyte formation.

The SCR reflects the proportion of schizonts within a given IDC that commit to gametocytogenesis and produce sexual ring stage progeny. Accurate calculation of SCRs hence relies on the simultaneous quantification of either asexually and sexually committed schizonts in the commitment cycle or of asexual parasites and early stage gametocytes in the immediate progeny, i.e. within the first 48 hours after invasion. However, since gametocytes are morphologically indistinguishable from asexual parasites until day three of sexual differentiation (stage II), the reliable determination of SCRs has traditionally been very laborious and technically challenging. Several laboratories have therefore developed flow cytometry-based assays that allow for an accurate measurement of gametocytemia in parasites expressing fluorophores or fluorophore-tagged gametocyte markers under the control of ectopic gametocyte-specific promoters (22, 23, 37, 40, 45). These assays enable distinguishing gametocytes from asexual stage parasites before they become morphologically distinct and therefore allow minimizing effects that may confound a precise determination of SCRs. Potential confounding factors include (i) the erroneous counting of gametocytes originating from multiple previous IDCs, (ii) the effect of multiplying asexual parasites and, in case of probing drugs or drug-like molecules, (iii) the potential lack of activity of compounds on early gametocyte survival. Because of the reduced sensitivity of gametocytes to most antimalarials (46, 47), it is important to identify and exclude confounding effects emerging from differential survival between asexual and sexual stage parasites. Hence, compared to standard light microscopy-based setups, flow cytometry-based assays greatly improved the accuracy and likewise the throughput of measuring SCRs in parasite populations (22, 23, 37, 40, 45).

Here, we developed a novel high content imaging (HCI) assay for the precise quantification of sexual commitment rates in *P. falciparum* parasites. This assay identifies sexually committed ring stage parasites based on the expression of endogenous mScarlet-tagged AP2-G, the earliest and most specific marker for sexually committed ring stages. Given the potential impact of antimalarials on malaria transmission, we used this assay to test a comprehensive panel of drugs for their possible effects on sexual commitment. In addition, we also included a diverse collection of other therapeutics that are commonly prescribed in malaria-endemic regions, including antihelminthics and analgesics, to account for a possible role of general stress-inducing factors on the sexual commitment process. Our results provide a systematic evaluation of the links between drug treatment and sexual commitment and suggest that antimalarial drug treatment does not promote transmission stage formation in *P. falciparum*.

## Materials and Methods

### Parasite culture

Intra-erythrocytic *P. falciparum* stages were cultured and synchronized as described (48, 49). Generally, parasites were grown in AB+ or B+ human RBCs (Blood Donation Center, Zürich, Switzerland) at a hematocrit of 5% in parasite culture medium consisting of 10.44 g/L RPMI-1640, 25 mM HEPES, 100 μM hypoxanthine, 24 mM sodium bicarbonate and 0.5% AlbuMAX II (Gibco #11021-037). The medium was further complemented with 2 mM choline chloride (Sigma #C7527) to maintain low background sexual commitment rates as observed in the presence of human serum (10). Cultures were gassed with 3% O_2_, 4% CO_2_ and 93% N_2_ and incubated in an airtight incubation chamber at 37°C.

### Cloning of transfection constructs

CRISPR/Cas9-based genome engineering of the NF54/ap2g-mScarlet, TM90C2B/ap2g-mScarlet and NF54/ap2g-re9h parasites was performed using a two-plasmid approach as previously described (10, 16). This system is based on co-transfection of a suicide and a donor plasmid. The suicide plasmid contains the expression cassettes for the Cas9 enzyme, the single guide RNA (sgRNA) and the human dihydrofolate reductase (hDHFR) resistance marker (pH-gC). A pD-derived donor plasmid was used for homology-directed repair of the Cas9-induced DNA double strand break (16).

The pH-gC_ap2g-3’ suicide plasmid targeting the 3’ end of *pfap2-g* has been described previously (10). The donor plasmid pD_ap2g-mScarlet was generated by assembling (i) the BamHI and SfoI-digested pD_ap2g-gfp plasmid (10) with a PCR product containing (ii) the *mScarlet* sequence preceded by nucleotides encoding a GSAG linker using the primers mScarlet-F and mScarlet-R amplified from a *P. falciparum* codon-optimized synthetic *mScarlet* sequence (50), and (iii) the 3’ homology region amplified from the pD_ap2g-gfp plasmid (10) using the primers ap2-g-3’-F and ap2-g-3'-SfoI-R in a Gibson reaction (51). The donor plasmid pD_ap2g-re9h was generated by assembling two PCR products using (1) the primers iso_re9h-F and iso_re9h-R amplified from pD_ap2g-mScarlet, and (52) re9h-F and re9h-R to amplify the *re9h* fragment (53) from pTRIX2-re9h (54). The sequence of the self-cleaving peptide T2A was included in the primers iso_re9h-R and re9h-F (55). Primer sequences used for cloning are listed in Table S1.

### Transfection and selection of gene edited parasites

*P. falciparum* transfection using the CRISPR/Cas9 suicide and donor plasmid approach was performed as described previously (16). Briefly, 50 μg of each of the suicide plasmid (pH-gC_ap2g-3’) and the respective donor plasmid (pD_ap2g-mScarlet or pD_ap2g-re9h) were co-transfected. Transgenic parasites were selected with 4 nM WR 99210 24 h after transfection for 6 days. Transgenic populations were usually obtained 2–3 weeks after transfection and correct editing of the *ap2-g* locus was then confirmed by PCR on gDNA. Primer sequences used for these PCRs are listed in Table S1.

### Drug assays

The screening for sexual commitment-inducing compounds was performed in a 96-well plate format and compounds were tested in twelve concentrations using two-step or three-step serial dilutions. Stock solutions of 10 mM drug were prepared in DMSO, except for chloroquine, acetaminophen, Aspirin, diclofenac, ibuprofen and piparaquine (prepared in -SerM). Working solutions were prepared in -SerM medium immediately before the experiment and 100 µL each was dispensed into the wells of a cell culture plate (Corning Incorporated, 96-well cell culture plate, flat bottom, REF 3596). Synchronous asexual parasites at 20-26 hpi and a parasitaemia of 0.5-1% were washed and resuspended in -SerM medium complemented with 10 mM choline chloride at 2. 5 % hematocrit. 100 µL parasite suspension was then added to each well and gently mixed with the compounds. As a positive control for sexual commitment-inducing conditions, parasites resuspended in -SerM medium lacking choline chloride were used. Plates were gassed and incubated in an airtight incubation chamber at 37°C for 48 hours.

### Quantification of parasite survival

At 20-26 hpi after reinvasion into new RBCs (i.e. 48 hours after the start of the assay), the parasitaemia was determined using flow cytometry. To this end, 40 μl parasite suspension was transferred from each well to a new 96-well plate (Corning Incorporated, 96-well cell culture plate, round bottom, REF 3788) and the samples were stained for 20 min with 40 μl 2X SYBR Green DNA stain (Invitrogen S7563) and then washed twice in 200 μl PBS. Plates were spun at 280 g for 2 min in between each step. To determine the parasitaemia, 200,000 events per sample were measured using the MACS Quant Analyzer 10. Data was analysed using the FlowJo_v10.6.1 software. The gating strategy removed small debris and doublets (two cells per measurement), and iRBCs were distinguished from uninfected RBCs based on the SYBR Green intensity (Fig. S6). Mean survival rates were calculated relative to -SerM/choline controls (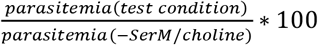) from three independent biological replicates. Curve fitting was performed using non-linear, four parameter regression model with variable slope (Graph Pad Prism, version 8.2.1).

### Quantification of sexual commitment rates by high content imaging

The SCR is defined as the proportion of sexually committed parasites in the total population. High content imaging (HCI) microscopy and automated image analysis were used to detect the number of all parasites based on DNA staining alone, whereas the sexually committed parasites were recognized via both DNA staining and AP2-G-mScarlet fluorescence. At 20-26 hpi after reinvasion into new RBCs (i.e. 48 hours after the start of the assay), cultures were stained with Hoechst (2.5 μg/mL) for 20 min and washed twice in 200 μL PBS. Plates were spun at 280 g for 2 min in between each step. The cultures were then diluted in PBS to a hematocrit of 0.075% and 200 μL were transferred to a clear-bottom 96-well HCI plate (Greiner CELLCOAT microplate 655948, Poly-D-Lysine, flat μClear bottom). Cells were allowed to settle for 15 min before image acquisition with an ImageXpress Micro widefield high content screening system (Molecular Devices) in combination with the MetaXpress software (version 6.5.4.532, Molecular Devices) and a Sola SE solid state white light engine (Lumencor). Filtersets for Hoechst (Ex: 377/50 nm, Em: 447/60 nm) and mScarlet (Ex: 543/22 nm, Em: 593/40 nm) were used with exposure times of 80 ms and 600 ms, respectively. 36 sites per well were imaged using a Plan-Apochromat 40x objective (Molecular Devices, cat# 1-6300-0297). Automated image analysis was performed using the MetaXpress software (version 6.5.4.532, Molecular Devices). Hoechst-positive as well as mScarlet-positive parasites were identified using a modular image analysis workflow built within the ImageXpress software (Molecular Devices) described in table S2, allowing for the calculation of sexual commitment rates (i.e. the proportion of Hoechst/mScarlet double-positive cells among all Hoechst-positive cells). SCRs were calculated as follows 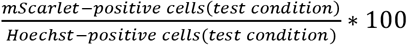. Mean relative SCRs were calculated relative to -SerM controls 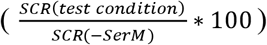 from three independent biological replicates.

### Quantification of RE9H luminescence intensity

At 20-26 hpi after reinvasion into new RBCs (i.e. 48 hours after the start of the assay), 180 μL of NF54/ap2g-re9h parasite culture was transferred from each well to a black 96-well plate (Greiner CELLSTAR microplate 655086, F-bottom, black) and incubated with 20 μL D-Luciferin (3.75 mg/mL in PBS) (Perkin Elmer, catalog# 122799) for 5-10 min at room temperature. Luminescence intensities were then measured using an intra-vital imaging system (Perkin Elmer, Lumina II) by exposing the plates for 3 min. Luminescence counts per well were determined by fitting a grid over the plate using the software Living Image (version 4.7.2).

### Quantification of gametocyte production rates by light microscopy

At 20-26 hpi, parasites were exposed to either -SerM or -SerM/choline conditions, gassed and incubated in an airtight incubation chamber at 37°C for 48 hours to complete the IDC and produce ring stage progeny. At this time point, parasitaemia was determined using Giemsa-stained blood smears (day 1). The culture medium was replaced with culture medium containing 50 mM N-acetyl-D-glucosamine to eliminate asexual parasites and the cultures incubated for 72 hours with daily medium changes (28). Gametocytaemia was determined by Giemsa-stained blood smears on day 4 (stage II gametocytes). Sexual commitment rates were calculated by dividing the gametocytemia determined on day 4 by the total parasitaemia determined on day 1 using results obtained from three independent biological replicates.

## Results

### An assay to quantify sexual commitment rates in *P. falciparum*

De-repression of the *ap2-g* locus marks the earliest known transcriptional event of sexual commitment. Here, we used a CRISPR/Cas9 gene editing strategy to fuse the *ap2-g* gene in frame to a sequence coding for the red fluorescent protein mScarlet (Figs. 1A and S1A,B). Using these NF54/ap2g-mScarlet parasites, we established an HCI-based assay to quantify the SCRs in live parasite populations via monitoring expression of the fluorescently tagged AP2-G-mScarlet reporter protein (Fig. S1C). In brief, the sexual commitment assay is initiated by seeding highly synchronous NF54/ap2g-mScarlet parasites in 96-well flat-bottom cell culture plates at the late ring/early trophozoite stage (20-26 hpi) at 1.25% haematocrit and 0.5-1% parasitaemia and exposing them to test conditions. After a 48-hour incubation period, i.e. during early intra-erythrocytic development of the subsequent generation, parasites are stained using the DNA dye Hoechst and transferred to a 96-well imaging plate at a haematocrit of 0.075%. The proportion of sexually committed ring stages/early stage I gametocytes (AP2-G-mScarlet-positive) among all iRBCs (Hoechst-positive) is quantified by fluorescence HCI microscopy. Imaging of 36 sites per well allows capturing information for 5000-10`0000 iRBCs. (Fig. 1B). At the same time point, parasitaemia is determined for each well of the cell culture plate in parallel by flow cytometry of SYBR Green-stained iRBCs to assess potential effects of the test conditions on parasite survival (multiplication).

**Figure 1.**
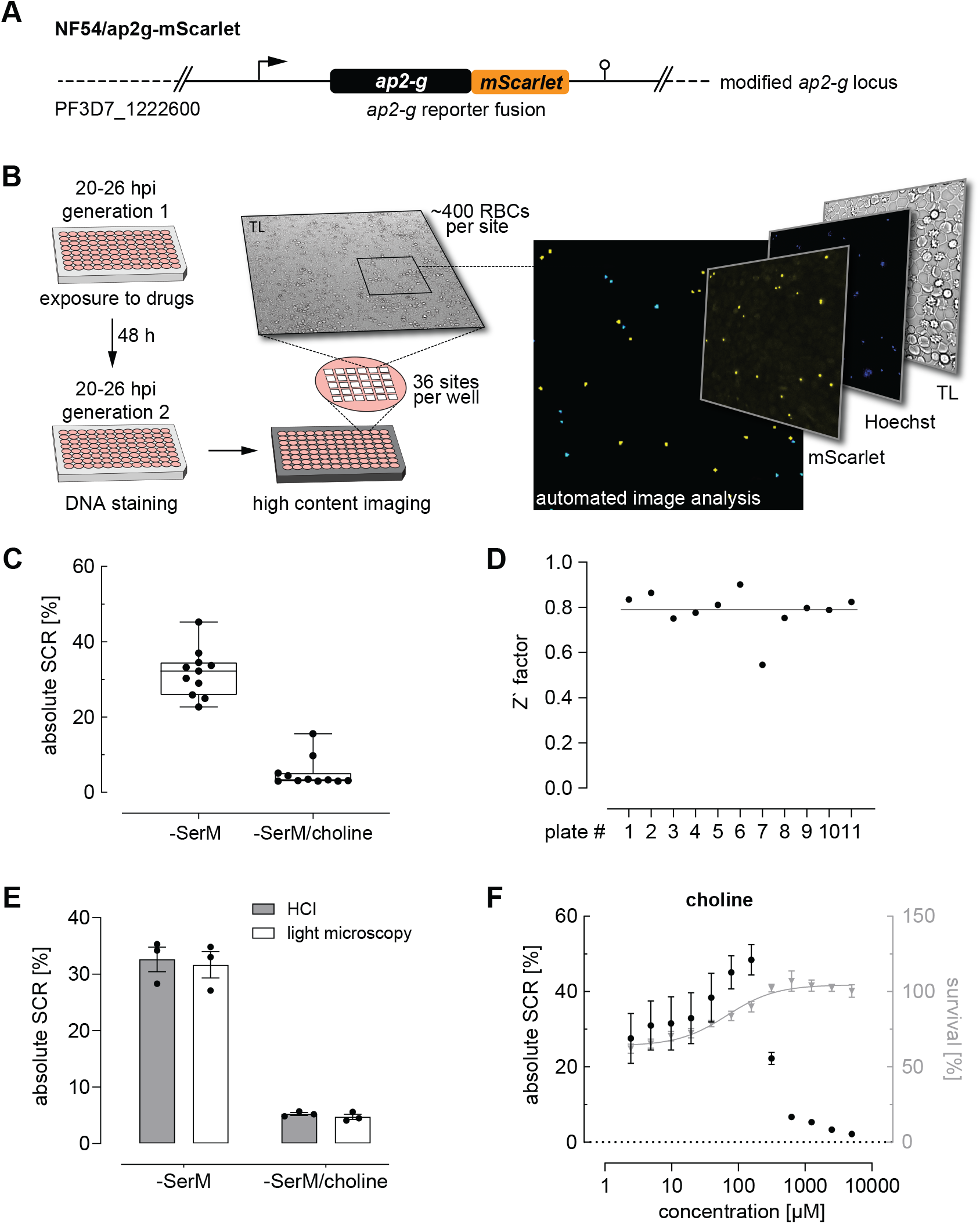
High content imaging-based quantification of SCRs. **(A)** Schematic of the modified *ap2-g* locus in NF54/ap2g-mScarlet parasites. mScarlet, red fluorescent protein. **(B)** Experimental setup of the high content imaging-based sexual commitment assay. TL, transmitted light. **(C)** Rates and variation of **s**exual commitment in NF54/ap2g-mScarlet parasites under SCR-inducing (-SerM) and inhibiting (-SerM/choline) control conditions. Boxplots show interquartile ranges; whiskers mark minimal and maximal values. Black bullets represent the mean SCRs measured in individual wells (obtained from 6 wells per screening plate) for eleven independent biological replicate experiments **(D)** Z′ factors determined for each screening plate. n=11; the line indicates the mean SCR across all plates. **(E)** SCRs and gametocyte formation rates under SCR-inducing (-SerM) and inhibiting (-SerM/choline) conditions quantified by high content imaging (HCI) and by light microscopy of stage II gametocytes on Giemsa-stained blood smears. n=3, error bars represent the standard error of the mean. **(F)** Dose-response effect of choline on the SCR (black bullets) and parasite survival (grey triangles). Data points represent the mean of three independent biological replicate experiments. Error bars represent the standard error of the mean.

To establish and validate this HCI assay, we used parasites cultured in serum-free medium (-SerM) either in presence or absence of choline, the commitment-inhibiting metabolite of lysoPC (10). As expected, these +/− choline control conditions (-SerM/choline; -SerM) resulted in consistently low and high SCRs, respectively (Fig. 1C), and revealed high assay robustness throughout the experiments presented here (Z’-factor of 0.79 +/− 0.09, Fig. 1D). Importantly, SCRs determined by this HCI assay are highly consistent with the corresponding gametocyte formation rates observed by light microscopy of Giemsa-stained blood smears prepared three days after the HCI-based readout (Fig. 1E). In absolute numbers, the NF54/ap2g-mScarlet line showed a mean SCR of 31.7% (95% CI: 27.5-36.9%) under inducing conditions (-SerM) and a baseline SCR of 5.2% (95% CI: 2.5-7.6%) under commitment-repressing conditions (-SerM/choline). To further validate assay performance, we performed choline titration experiments that revealed half-maximal inhibition of sexual commitment at 275 μM choline (95% CI: 256-293 μM) and impaired parasite multiplication at low choline concentrations, which is consistent with previously published data (10) (Fig. 1F).

In summary, these data show that this HCI-based assay allows capturing SCRs under controlled conditions and at a throughput that facilitates the systematic and robust investigation of modulators of parasite sexual commitment.

### Many antimalarials induce sexual commitment at growth-limiting concentrations

We used the above assay to investigate a total of 29 pharmaceuticals for potential effects on sexual commitment in 12-point dose-response assays. In addition to 14 antimalarials, we also screened 15 drugs commonly used in malaria endemic regions, such as analgesics and antihelminthics (Table 1). Compounds showing SCR-inducing effects during primary screening were validated using three independent biological replicate experiments.

**Table 1.** Compounds tested for activities on *P. falciparum* sexual commitment. Studies that observed an elevating effect of antimalarials on gametocyte production are indicated. S/P, sulfadoxine/pyrimethamine (“Fansidar”).

| **Antimalarials** | *effect on SCR reported in:* |
| --- | --- |
| amodiaquine |  |
| atebrin |  |
| atovaquone | (37) |
| chloroquine | (35-37, 81) |
| dihydroartemisinin | (38) |
| lumefantrine |  |
| mefloquine | (37) |
| piperaquine | (37) |
| primaquine | (37) |
| proguanil |  |
| pyrimethamine | (39) |
| pyronaridine |  |
| quinine |  |
| sulfadoxine | (39) |
| **Antihelminthics** |  |
| albendazole |  |
| ivermectin |  |
| mebendazole |  |
| moxidectin |  |
| praziquantel |  |
| **Antipyretics** |  |
| acetylsalicylic acid |  |
| ibuprofen |  |
| acetaminophen |  |
| diclofenac |  |
| **Antibiotics** |  |
| azithromycin |  |
| doxycycline |  |
| **Antidiabetics** |  |
| gliquidone |  |
| insulin |  |
| metformin |  |
| **Steroids** |  |
| dexamethasone |  |

While most antimalarials showed a trend towards elevating parasite SCR, this effect was restricted to a narrow concentration window and generally remained linked to drug levels that inhibited asexual parasite replication (Figs. 2A and S2). Chloroquine, pyrimethamine and mefloquine, for instance, had no prominent effect at low concentrations but elevated the parasite SCR just below the IC50 (Fig. 2B). At higher drug concentrations, SCRs fluctuated markedly and reached high values in some instances. However, as these elevated SCRs were observed in populations with complete or near-complete inhibition of parasite survival, they did not result in an enhanced production of sexually committed parasites. In fact, when placed in the context of parasite survival, only specific sub-IC50 concentrations of the widely used mefloquine and pyrimethamine led to a significant increase in sexual ring stage formation on the absolute scale (Fig. 2C). While the effect of pyrimethamine was minor, mefloquine, which is known to inhibit protein synthesis via direct binding to 80S ribosomes (56), increased the formation of ring stage gametocytes by a factor of 2.4 (95% CI: 1.2 to 4.7) at a concentration of 3.9 nM, i.e. several magnitudes below therapeutic concentrations (52, 57).

**Figure 2.**
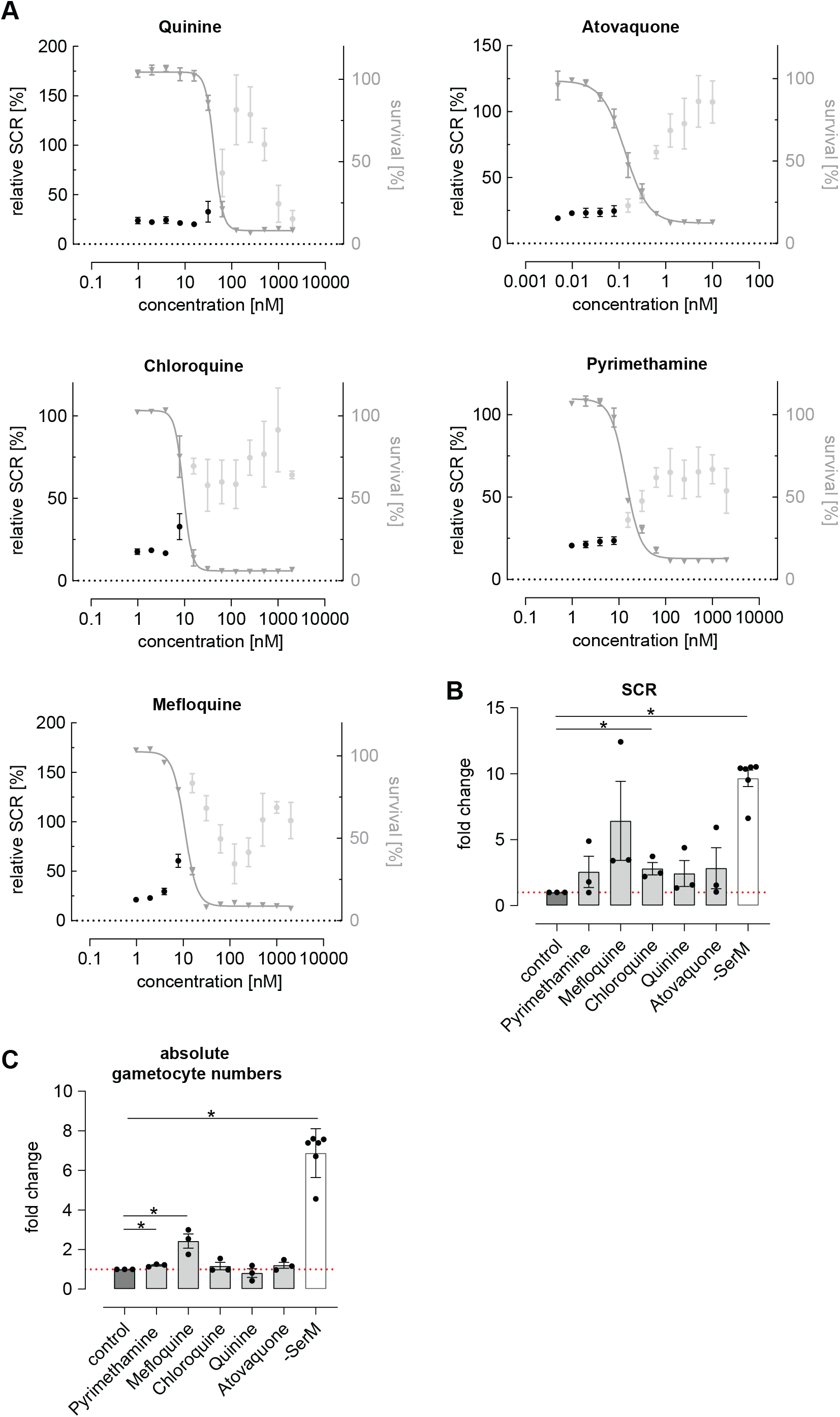
Dose-response relationship between antimalarial compounds and parasite sexual commitment. **(A)** Antimalarials show a general trend towards induction of sexual commitment at growth-inhibiting concentrations. Mean parasite survival rates and SCRs are indicated by grey triangles and black bullets, respectively. Grey bullets represent SCRs at compound concentrations above the IC50. Values are normalized to the corresponding control conditions (-SerM for SCR) and (-SerM/choline for survival). Data points represent the mean of three independent biological replicate experiments. Error bars represent standard error of the mean. **(B)** Maximal SCR increase observed in response to sub-IC50 drug levels. Bars indicate mean fold changes of SCRs compared to control conditions (untreated, -SerM/choline/DMSO), with black bullets representing fold changes from individual biological replicates. Fold changes are defined as 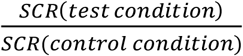. For each drug, the results shown were derived from the concentration for which the maximal net increase in absolute sexual ring stage formation was observed (pyrimethamine: 7.8 nM; mefloquine: 7.8 nM; chloroquine: 7.8 nM; quinine: 31.25 nM; atovaquone: 0.01 nM). -SerM shows the effect on SCRs obtained upon choline depletion. Asterisks mark significant differences (p-value < 0.05; paired two-tailed Student's t-test). n=3; error bars represent the standard error of the mean. **(C)** Exposure to sub-therapeutic mefloquine concentrations result in an absolute increase of sexually committed ring stage progeny formed. Pyrimethamine shows a similar but markedly less pronounced effect. Bars indicate mean fold changes of sexual ring stage formation compared to untreated control conditions (-SerM/choline/DMSO), with black bullets representing fold changes from individual biological replicates. Fold changes are defined as 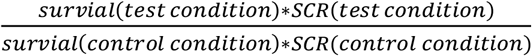. For each drug, the results shown were derived from the concentration for which the maximal net increase in absolute sexual ring stage formation was observed (pyrimethamine: 3.9 nM; mefloquine: 3.9 nM; chloroquine: 7.8 nM; quinine: 3.9 nM; atovaquone: 0.01 nM). -SerM shows the effect on SCRs obtained upon choline depletion. Asterisks mark significant differences (p-value < 0.05; paired two-tailed Student's t-test). n=3; error bars represent the standard error of the mean.

Noteworthy, parasites treated with artemisinin or its derivatives dihydroartemisinin (DHA), artemether and artesunate, emitted autofluorescence at various wavelengths including the TRITC channel, making a fluorescence-based quantification of SCRs impossible. To circumvent this issue and evaluate the effect of DHA, for which a sexual commitment-inducing effect has recently been demonstrated (38), we generated the NF54/ap2g-re9h reporter line (Fig. 3A and Fig. S3). These parasites express an AP2-G-T2A-RE9H luciferase fusion protein from the endogenous *ap2-g* locus and hence allow using luminescence as a proxy for quantifying AP2-G expression (Fig. 3B). In contrast to the fluorescence-based assay described above, the NF54/ap2g-re9h line enables determining absolute levels of sexual ring stages formed in parasite populations rather than quantifying SCRs at a single cell level. The NF54/ap2g-re9h cell line showed robustness in reporting sexual commitment under control conditions (Z’-factor of 0.57 +/−0.13) as well as the ability to capture the dose-dependent effect of choline on SCRs (half-maximal inhibition of sexual commitment at 194 μM; 95% CI: 136-298 μM) (Fig. 3C).

**Figure 3.**
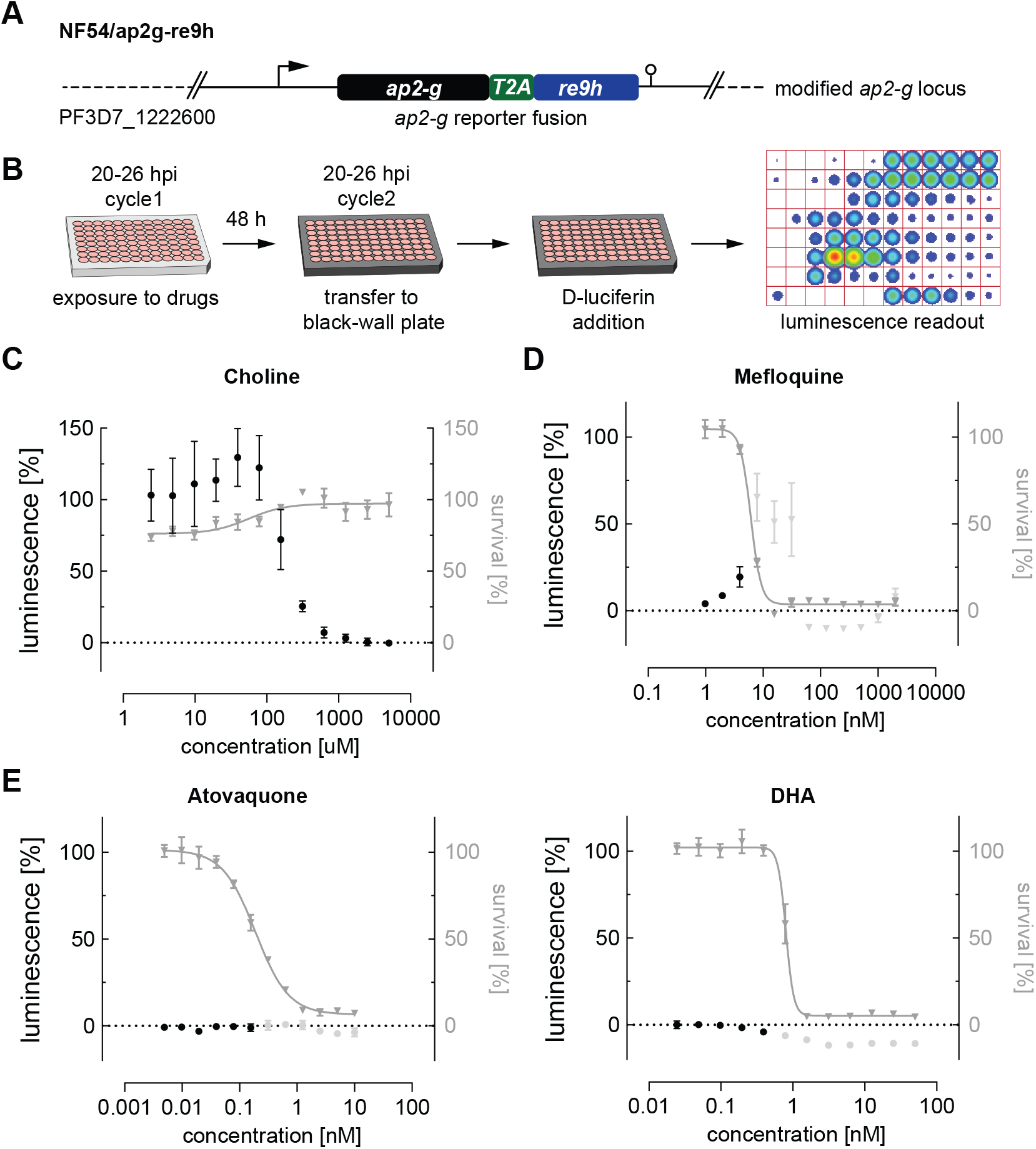
Luciferase-based quantification of sexual commitment. **(A)** Schematic of the modified *ap2-g* locus in NF54/ap2g-re9h parasites. *re9h*, gene encoding red-shifted firefly luciferase RE9H. T2A, self-cleaving peptide. **(B)** Experimental setup of RE9H luciferase-based quantification of SCRs. **(C)** Dose-response effect of choline on the SCR (black bullets) and parasite survival (grey triangles). Data points represent the mean of three independent biological replicate experiments. Error bars represent the standard error of the mean. **(D)** Mefloquine induces sexual commitment within a narrow sub-therapeutic window. Mean parasite survival rates and relative RE9H reporter-mediated luminescence as a surrogate for SCRs are indicated by grey triangles and black bullets, respectively. Grey bullets represent SCRs at compound concentrations above the IC50. Values are normalized to the corresponding control conditions (-SerM for SCR) and (-SerM/choline for survival). n=3; error bars represent standard error of the mean. **(E)**Neither atovaquone nor DHA induce sexual commitment. Mean parasite survival rates and relative RE9H reporter-mediated luminescence as a surrogate for SCRs are indicated by grey triangles and black bullets, respectively. Grey bullets represent SCRs at compound concentrations above the IC50). Values are normalized to the corresponding control conditions (-SerM for SCR) and (-SerM/choline for survival). n=3; error bars represent standard error of the mean.

After having validated the use of NF54/ap2g-re9h parasites for screening purposes, we made use of this line to quantify the effect of selected drugs on sexual commitment. These experiments confirmed the SCR-inducing effect of mefloquine at sub-therapeutic conditions (Fig. 3D), and the lack thereof after treatment with Atovaquone (Fig. 3E), corroborating the results obtained with the NF54/ap2g-mScarlet line (Fig. 2A). Importantly, no increase in SCRs was observed following exposure to DHA (Fig. 3E).

### Commonly prescribed drugs have no relevant effect on sexual commitment

Next, we investigated the effect of a collection of frequently used drugs, including antihelminthics, antibiotics as well as compounds used to treat pain, fever and inflammation (Table 1). Similar to what we observed for most antimalarials, antihelminthic drugs, including albendazole, ivermectin and moxidectin, showed trends towards increasing sexual commitment in the NF54/ap2g-mScarlet line (Figs. 4A and S4). For most compounds, however, this effect was restricted to parasite growth-inhibiting drug concentrations that are substantially higher than the maximum serum levels observed in patients after standard treatment (58–63). Interestingly, moxidectin, a macrocyclic lactone known to be active against *Plasmodium berghei* mosquito stages (64), also prevented asexual parasite replication rather effectively (IC50 = 159 nM) (Fig. 4A). While moxidectin induced sexual commitment at concentrations near the IC50, no net increase in ring stage gametocytes was observed (Fig. S4). The tested antipyretics (aspirin, ibuprofen, diclofenac and acetaminophen), antibiotics (azithromycin and doxycycline), antidiabetics (metformin, gliquidone) as well as dexamethasone, a corticosteroid with anti-inflammatory and immunosuppressant properties, had no effects on parasite sexual commitment at medically relevant concentrations (65–68) (Figs. 4B and S4).

**Figure 4.**
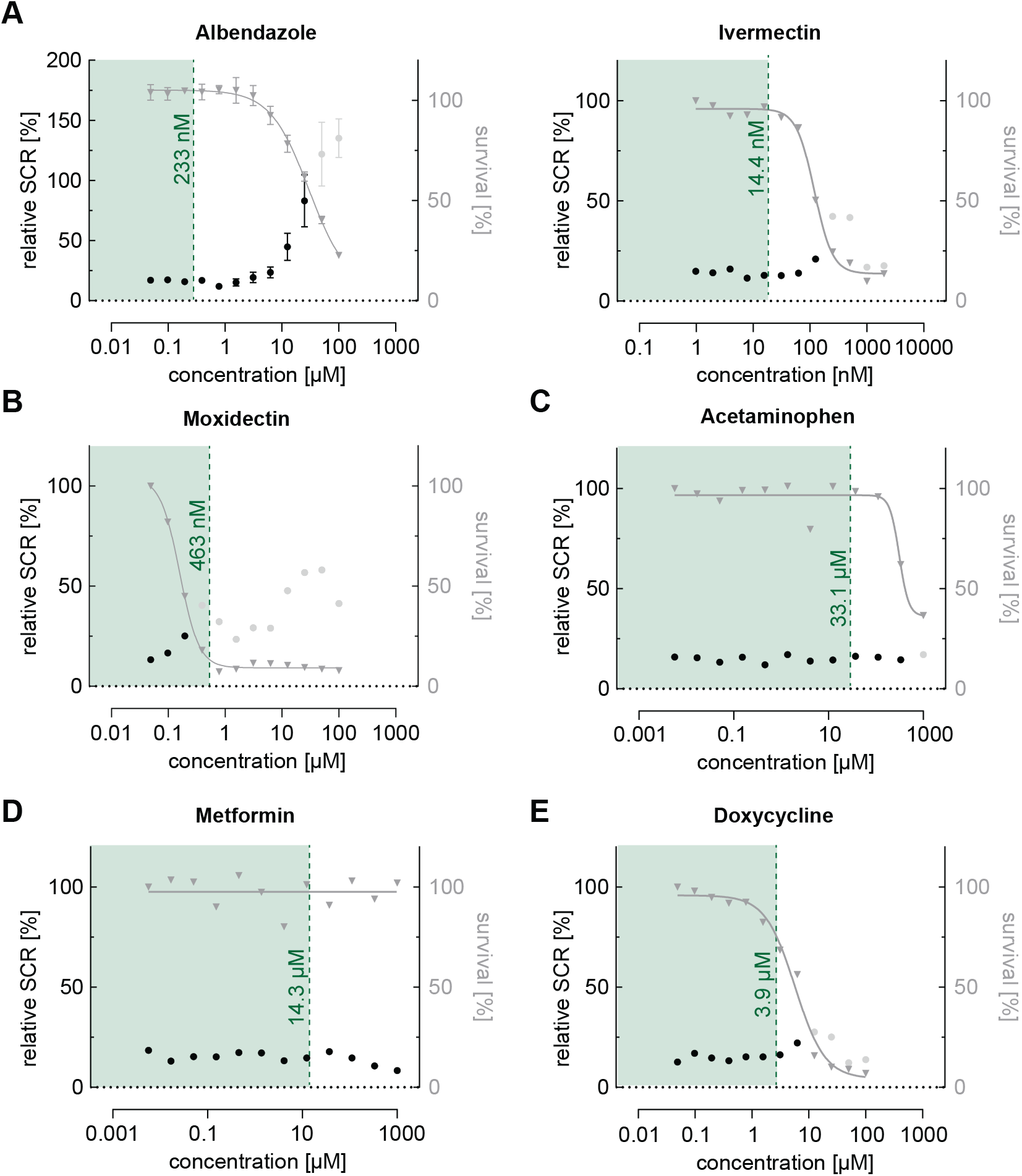
Dose-response relationship between commonly used drugs and parasite sexual commitment. **(A)** The antihelminthicsa albendazole and ivermectin show a general trend towards induction of sexual commitment at super-physiological concentrations that also affect parasite growth (see also Fig. S4). **(B)** The antihelminthic moxidectin affects parasite growth as well as SCRs at physiological concentrations. **(C)-(E)** The commonly used drugs acetaminophen (antipyretic) (C), metformin (antidiabetic) (D) and doxycycline (antibiotic) (E) have no effect on parasite growth or SCRs at physiologically relevant concentrations. Mean parasite survival rates and SCRs are indicated by grey triangles and black bullets, respectively. Grey bullets represent SCRs at compound concentrations above the IC50. Values are normalized to the corresponding control conditions (-SerM for SCR) and (-SerM/choline for survival). Albendazole: n=3; error bars represent standard error of the mean. Moxidectin, acetaminophen and metformin: n=1.

### Cellular stress modulates sexual commitment rather than target-specific activities

Drug resistance has been associated with increased gametocyte carriage in malaria patients with uncomplicated P. *falciparum* infections (41, 42, 69, 70). However, whether these observations are a consequence of a higher burden of asexually replicating parasites or higher rates of sexual commitment is unclear. To address this question, we tagged the *ap2-g* locus in the multidrug-resistant *P. falciparum* strain TM90C2B using the same CRISPR/Cas9 gene editing approach employed to generate the NF54/ap2g-mScarlet line (Figs. 5A and S5). The TM90C2B strain has reduced susceptibility to chloroquine, cycloguanil, pyrimethamine and atovaquone (71). As expected, transgenic parasites of the TM90C2B/ap2g-mScarlet cell line showed substantially higher tolerance to chloroquine and pyrimethamine compared to the non-resistant NF54/ap2g-mScarlet control line (Figs. 5B and 5C). For chloroquine, the IC50 increased approximately 8-fold from 9.4 nM (95% CI: 8.477 to 10.30 nM) to 71.2 nM (95% CI: 63.79 to 79.69 nM) and for pyrimethamine by a factor of 2`500 from 14.1 nM (95% CI: 12.79 to 15.42 nM) to 36.1 μM (95% CI: 22.9 to 175.9 μM). Similar to the observations made for NF54/ap2g-mScarlet parasites, chloroquine showed a dose-dependent trend towards increasing sexual commitment in the TM90C2B/ap2g-mScarlet line. Importantly, however, drug concentrations that led to increased SCRs in the drug-sensitive NF54/ap2g-mScarlet strain were ineffective in drug-resistant TM90C2B/ap2g-mScarlet parasites. In these multidrug-resistant parasites, increased SCRs were only observed at chloroquine concentrations that inhibited asexual replication beyond the IC50 (Fig. 5B). Likewise, pyrimethamine, for which we observed a minor but statistically significant inducing effect on the formation of sexual ring stages (see Fig. 2), elevated sexual commitment in TM90C2B/ap2g-mScarlet cells. Again, this activity was exclusively observed at drug concentrations that substantially inhibited parasite growth (>10 μM) and in this case even exceeded the serum concentrations observed in patients following standard treatment (median of 400 nM (72)).

**Figure 5.**
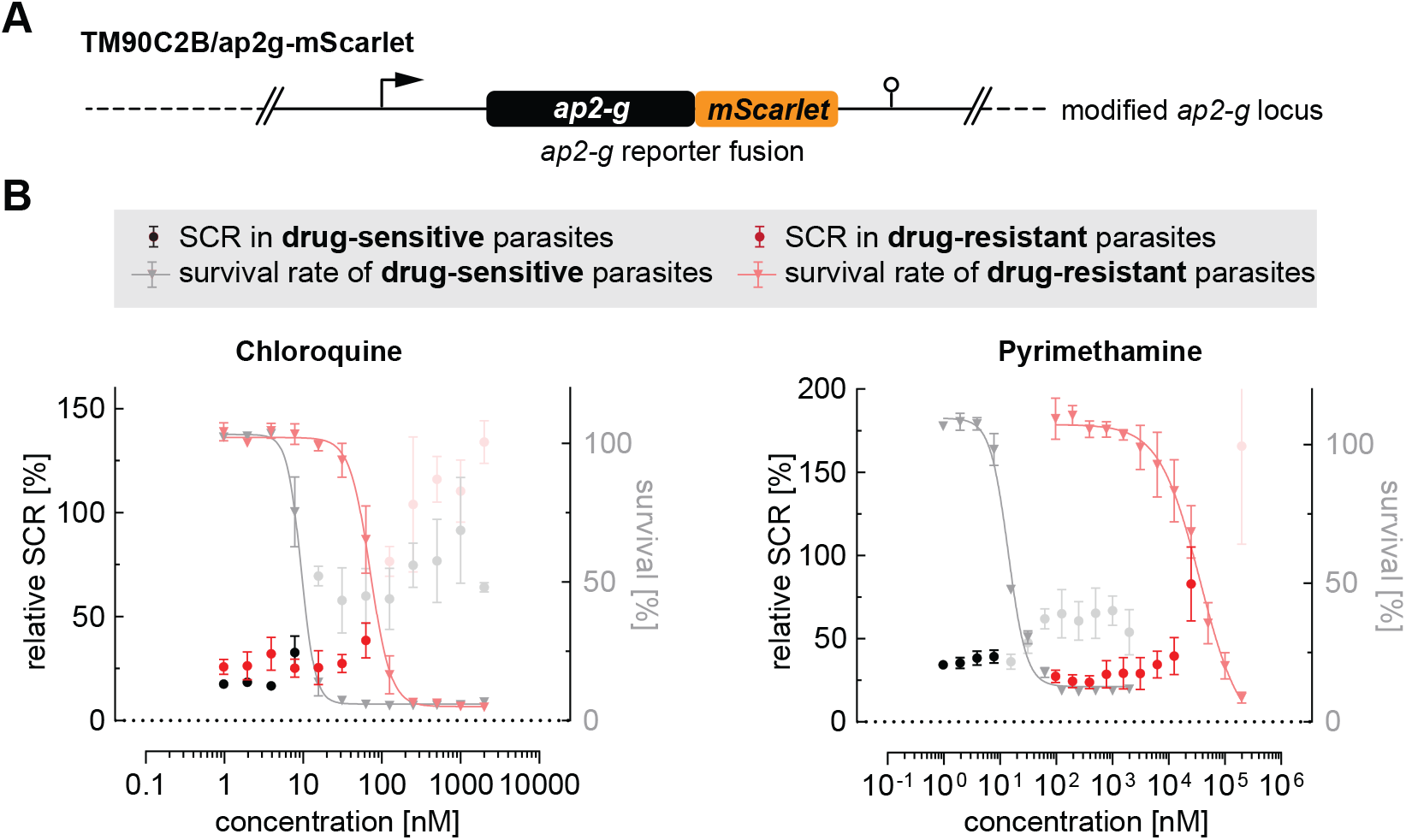
Effect of antimalarials on SCR in drug-resistant parasites. **(A)** Schematic of the modified *ap2-g* locus in multidrug-resistant TM90C2B/ap2g-mScarlet parasites. **(B)** Trends towards increased sexual commitment are linked to growth-inhibiting concentrations of Chloroquine and Pyrimethamine in drug-sensitive as well as in drug-resistant parasite lines. For NF54/ap2g-mScarlet, mean parasite survival rates and SCRs are indicated by grey triangles and black bullets, respectively (values adopted from Fig. 2A). For drug-resistant TM90C2B/ap2g-mScarlet parasites, mean parasite survival rates and SCRs are indicated by light red triangles and red bullets, respectively. Grey and light red bullets represent SCRs at compound concentrations above the IC50. Values are normalized to the corresponding control conditions (-SerM for SCR) and (-SerM/choline for survival). n=3; error bars represent standard error of the mean.

## Discussion

Malaria transmission relies on the formation of gametocytes from a pool of asexually replicating parasites. The proportion of parasites that differentiate into these sexual sages, the so-called sexual commitment rate, is variable and – at least to a certain extent – driven by cues in the microenvironment of intra-erythrocytic parasites (10, 23, 27–29, 37, 38). While a number of conditions, including exposure to antimalarial drugs, have been suggested or demonstrated to stimulate sexual commitment (10, 30–32, 34, 36–39, 73–75), the impact of therapeutic interventions on gametocyte production and malaria transmission is still a matter of debate. Differences in drug-susceptibility between *P. falciparum* asexual parasites and gametocytes, combined with the long period of sexual differentiation, render an evaluation of drug-induced effects on gametocytogenesis a challenging endeavor. Here, we established a high content imaging-based assay to systematically probe the impact of a comprehensive set of antimalarials and other drugs commonly prescribed in malaria-endemic regions on *P. falciparum* sexual commitment. Allowing for the simultaneous quantification of parasite survival and SCRs, this assay facilitated measuring net effects on gametocyte formation.

Consistent with previous observations, we found that *P. falciparum* parasites tend to elevate SCRs following *in vitro* exposure to a multitude of drugs, including, but not limited to antimalarials. While this effect was negligible for most tested compounds, some antimalarials markedly altered the rate at which parasites committed to the sexual pathway. Chloroquine for example, elevated SCR by a factor of 2.9 (95% CI: 1.4-6.1) at a concentration of 7.8 nM (see Fig. 2B) when compared to the untreated control population. This effect, however, was restricted to a narrow drug concentration window around the IC50 value and did not cause a net increase in sexual ring stages formed (see Fig. 2C).

Contrary to this general trend, exposure to pyrimethamine and mefloquine did not only elevate the SCR but also caused a low to moderate net increase in absolute numbers of sexual ring stages, respectively. These gametocytogenesis-promoting activities were linked to specific concentrations (3.9 nM for pyrimethamine; 3.9 nM for mefloquine) near the IC50 for both drugs. While the 2.4-fold increase in sexual ring stages formed (95% CI: 1.2-4.7) after exposure to 3.9 nM mefloquine was the highest activity observed throughout this study, this value is substantially lower compared to the gametocytogenesis-promoting effect observed under LysoPC/choline-depleted -SerM control conditions (fold change of 7.3; 95% CI: 6.1-8.7). Considering this relatively low drug-induced activity, as well as the narrow drug concentration window within which sexual commitment was elevated, it seems highly unlikely that drug treatment *per se* could have a relevant effect on promoting gametocyte production and malaria transmission in real life settings.

Nevertheless, our data reinforce the view that parasites can change rates of sexual commitment and probably also the absolute number of gametocytes formed in response to exposure to drugs at sub-curative levels. Fueled by previous reports about increased gametocytemias and mosquito infectivity following treatment of parasites with drug resistance mutations (42, 43), our observations thus raise the question as to whether therapeutic drug concentrations could provoke a disproportionally high rate of gametocyte formation in drug-resistant parasites. Using the multidrug-resistant parasite strain TM90C2B, we could not observe such effects for chloroquine and pyrimethamine and their activities on SCRs remained tightly linked to growth-inhibiting drugs levels. For instance, while we found the TM90C2B parasites to elevate SCRs in response to pyrimethamine exposure, this activity occurred only at a drug concentration of 25 μM, i.e. at growth-inhibiting concentrations close to the IC50 and >2′500-fold higher compared to the concentration that led to elevated SCRs in the pyrimethamine-sensitive NF54 strain.

The strict link between unfavorable growth conditions and elevated parasite SCRs strongly suggests that drug-induced sexual commitment is linked to general stress responses, rather than to compound-specific effects targeting the sexual commitment pathway. In fact, to date we are missing strong evidence for the ability of antimalarials or other drugs to interfere with the molecular process of variable gametocyte formation. Considering the involvement of epigenetic control mechanisms and phospholipid metabolism in the regulation of parasite sexual commitment (10–12), it would however not be surprising to observe corresponding effects for drugs interfering with these processes specifically. For instance, histone deacetylase inhibitors, which have important applications in anti-cancer treatments (76) and show promising activity against *P. falciparum* blood stage parasites (77, 78), may interfere with heterochromatin-mediated silencing of the *ap2-g* locus. Similarly, choline kinase inhibitors, for which a direct effect on parasite sexual commitment has previously been demonstrated (10), were proposed as new therapeutic tools against a variety of human diseases, including bacterial and parasitic infections (79). It will thus be important to carefully evaluate potential effects of such molecules on sexual commitment and gametocyte formation before developing them into antimalarial agents.

Based exclusively on *in vitro* experiments, it is clear that the data presented here cannot fully reflect the complex situation found in patients infected with *P. falciparum*. For example, the different microenvironments that parasites encounter at sequestration sites, including the bone marrow and spleen, may have profound effects on drug kinetics and bioavailability. Furthermore, in recent efforts, Portugaliza and colleagues simulated the short *in vivo* half-life of artemisinin/DHA by exposing *in vitro* cultured parasites to short drug pulses and identified stage-specific effects on sexual commitment (38). While ring stage populations exposed to 3-hour pulses of DHA responded with decreased SCRs, trophozoites showed elevated SCRs following drug pressure. By contrast, the experiments presented here did not reveal an effect of this artemisinin derivative on parasite SCR. These discrepancies are likely a result of the distinct experimental setups used – particularly the different periods of drug exposure used. Clearly, a comprehensive picture of physiologically relevant links between *P. falciparum* gametocyte production and drug pressure can only be gained by accounting for a variety of parameters including pharmacokinetics, pharmacodynamics as well as different host determinants and microenvironments. The limitations of our *in vitro* studies notwithstanding, the data presented here imply that none of the existing antimalarial drugs act specifically on the molecular pathways controlling sexual commitment and are hence unlikely to significantly enhance malaria transmission. This is in line with studies reporting that combination therapies, in particular ACTs, are associated with an effective reduction in gametocyte carriage (1, 2, 80) and indicates that the positive effect of antimalarial treatment clearly outweighs potential risks of increased transmission.

## Acknowledgements

We thank Christian Scheurer for providing the multidrug-resistant parasite line TM90C2B and Eilidh Carrington for supervising the engineering of the pD_ap2g-mScarlet plasmid. This work was supported by the Swiss National Science Foundation (grants 310030_200683 and BSCGI0_157729) and the Fondation Pasteur Suisse.

## Author Contributions

BT performed all experiments, analyzed, and interpreted the data. EH, AP and BT generated the transgenic parasite lines. TB performed experiments performed with NF54/ap2g-re9h parasites. NB and TV conceived of the study, designed and supervised experiments, and provided resources. NB prepared illustrations and wrote the manuscript, TV edited the manuscript. All authors contributed to the final editing of the manuscript.

---

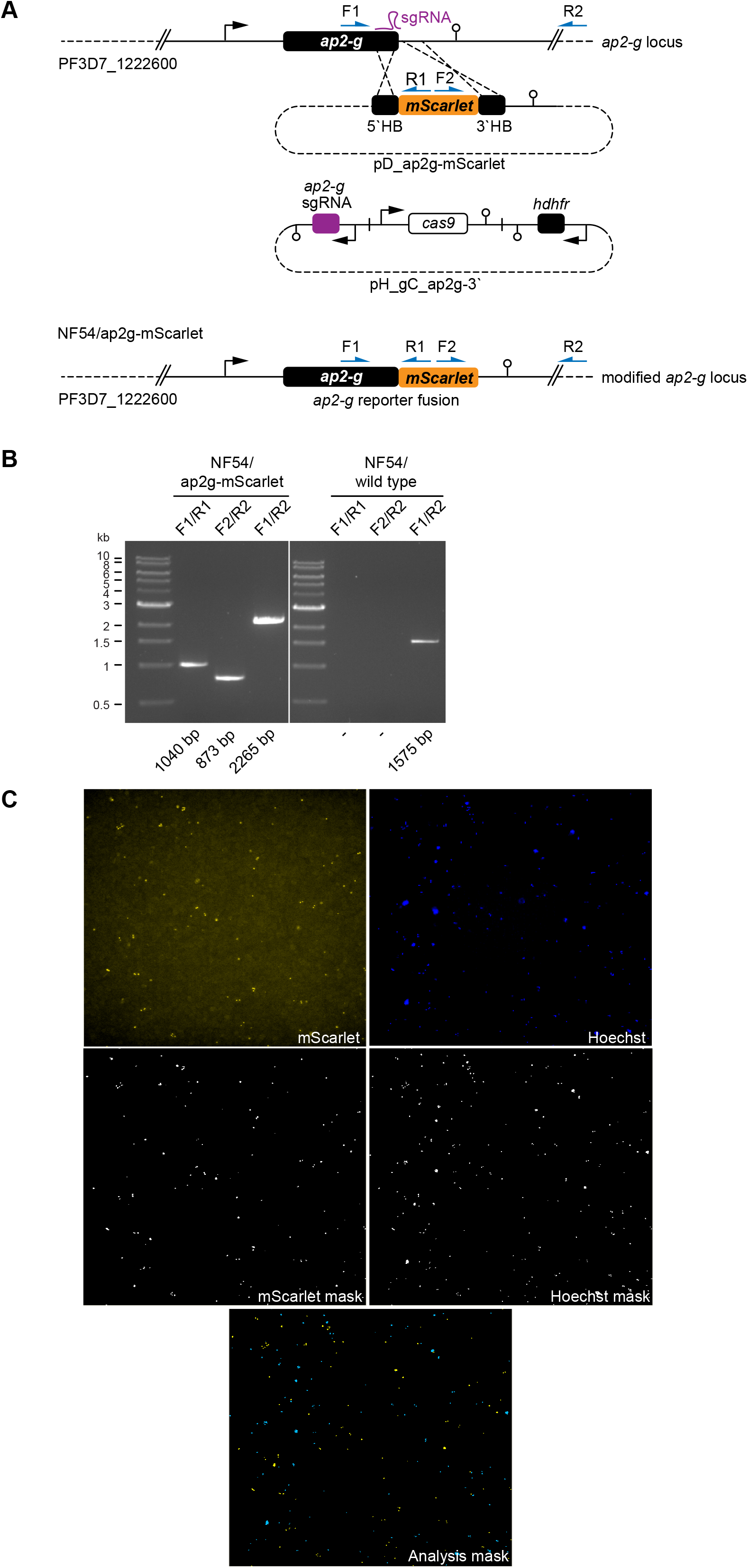

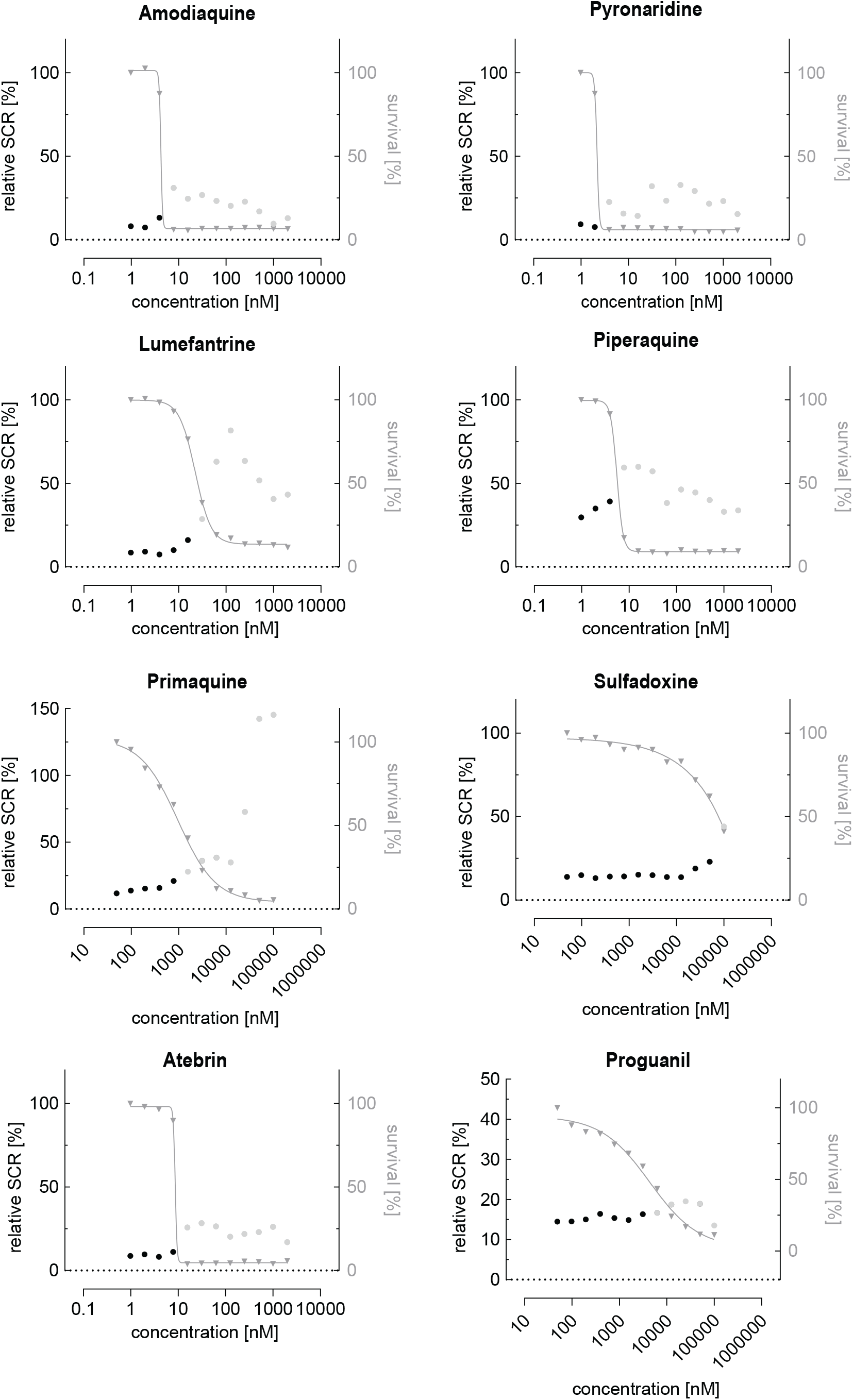

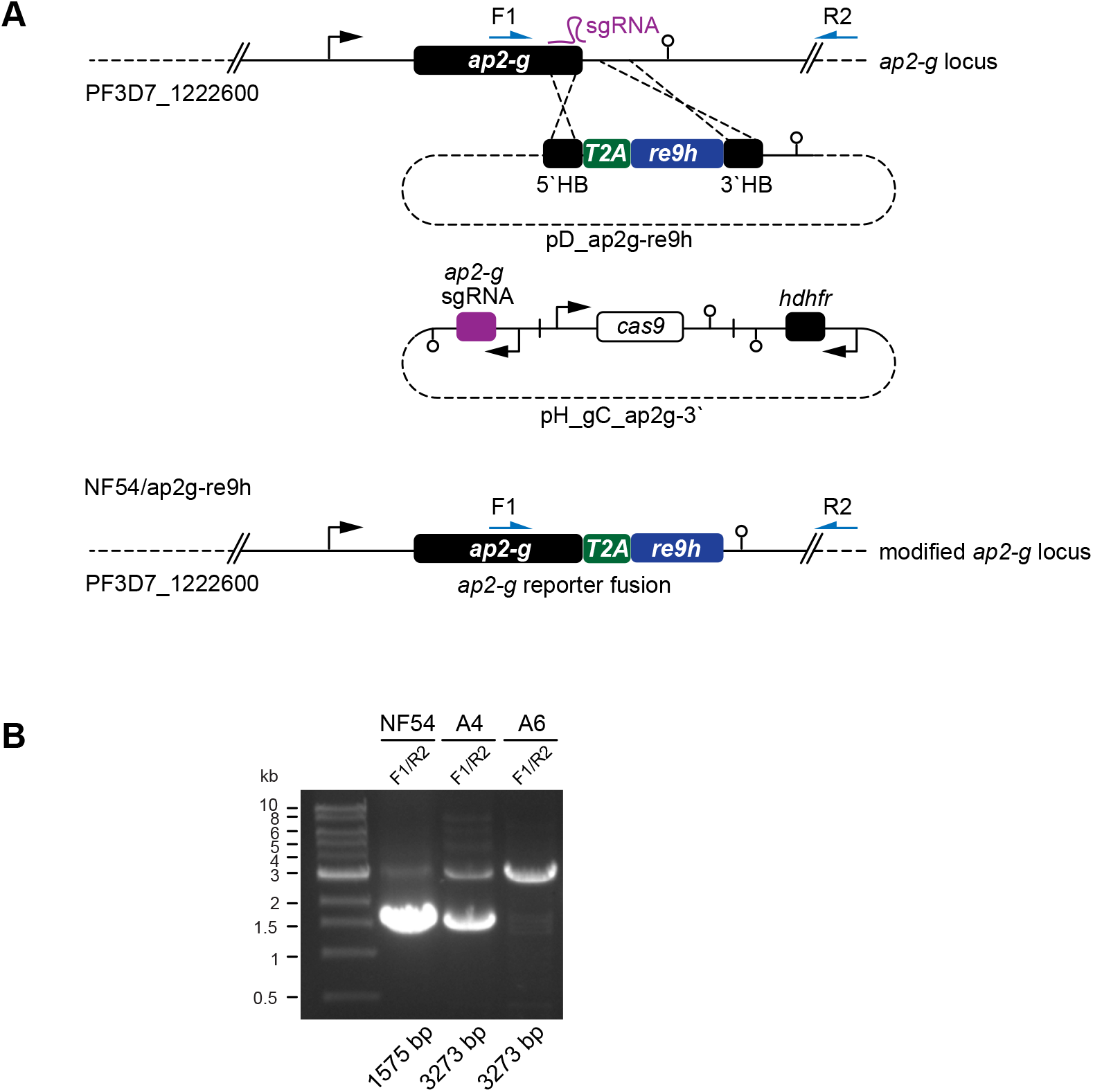

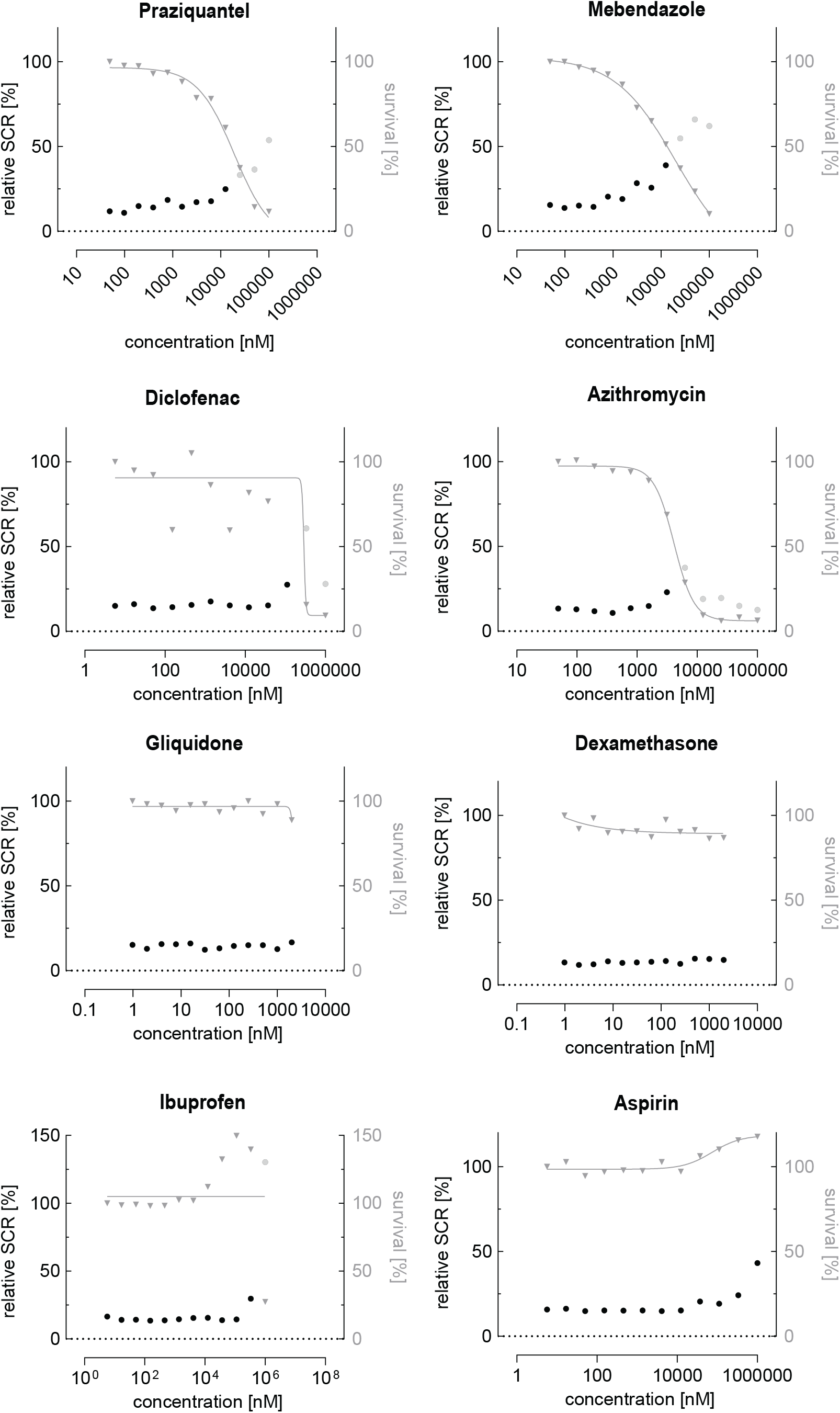

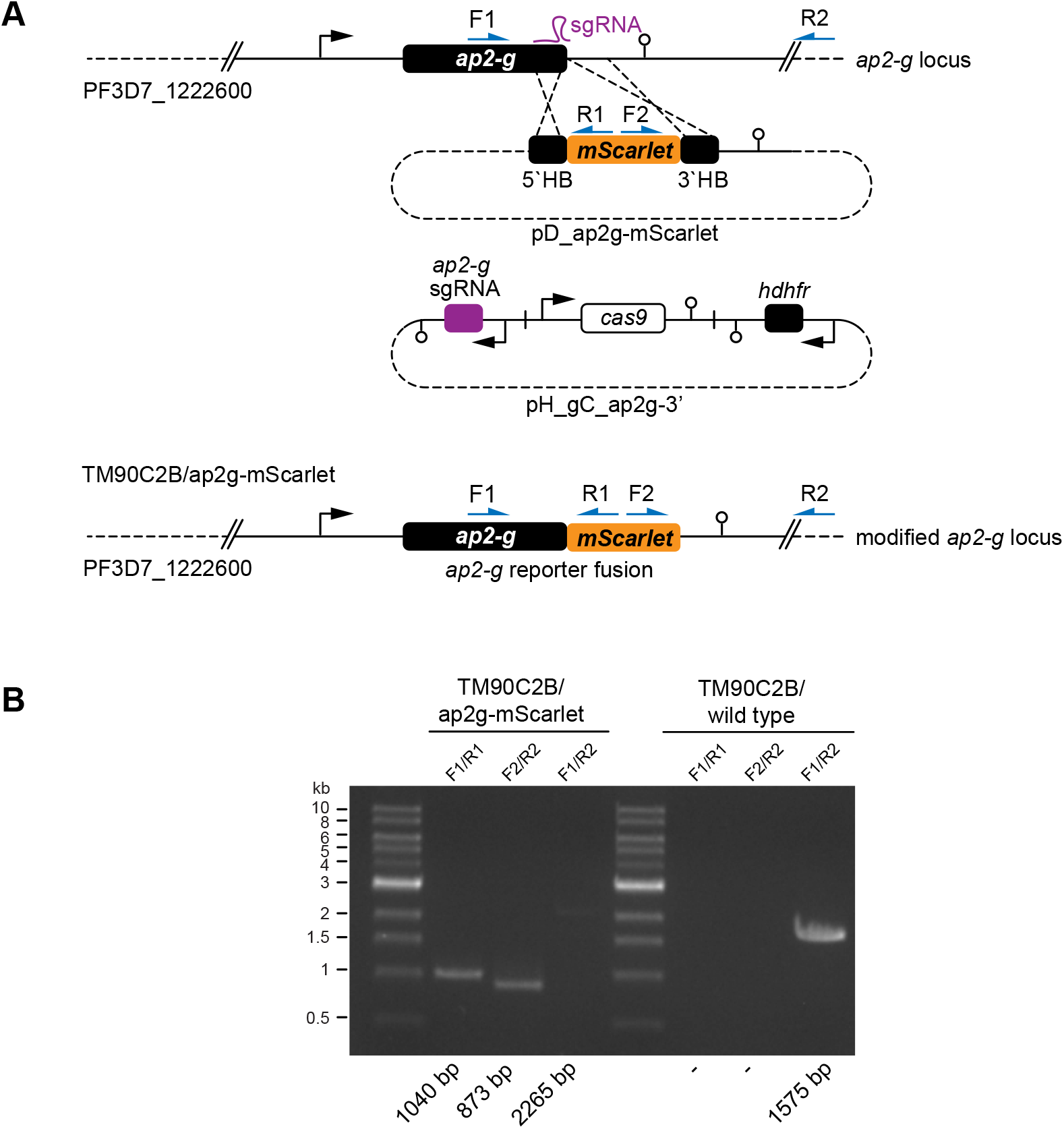

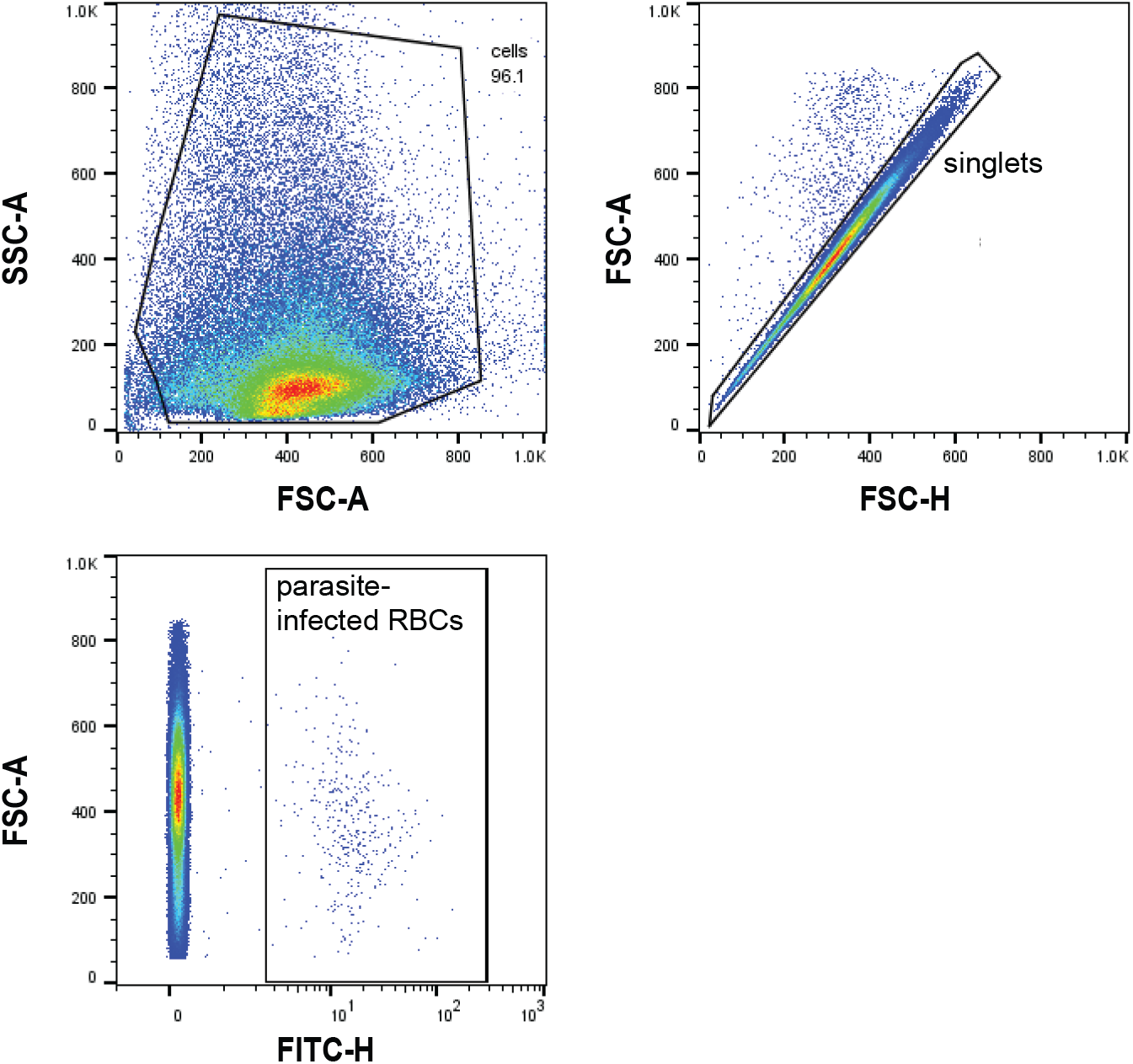

