## Supplemental Material for "Revisiting the effect of pharmaceuticals on transmission stage formation in the malaria parasite *Plasmodium falciparum*"

1 **Supplementary Material**

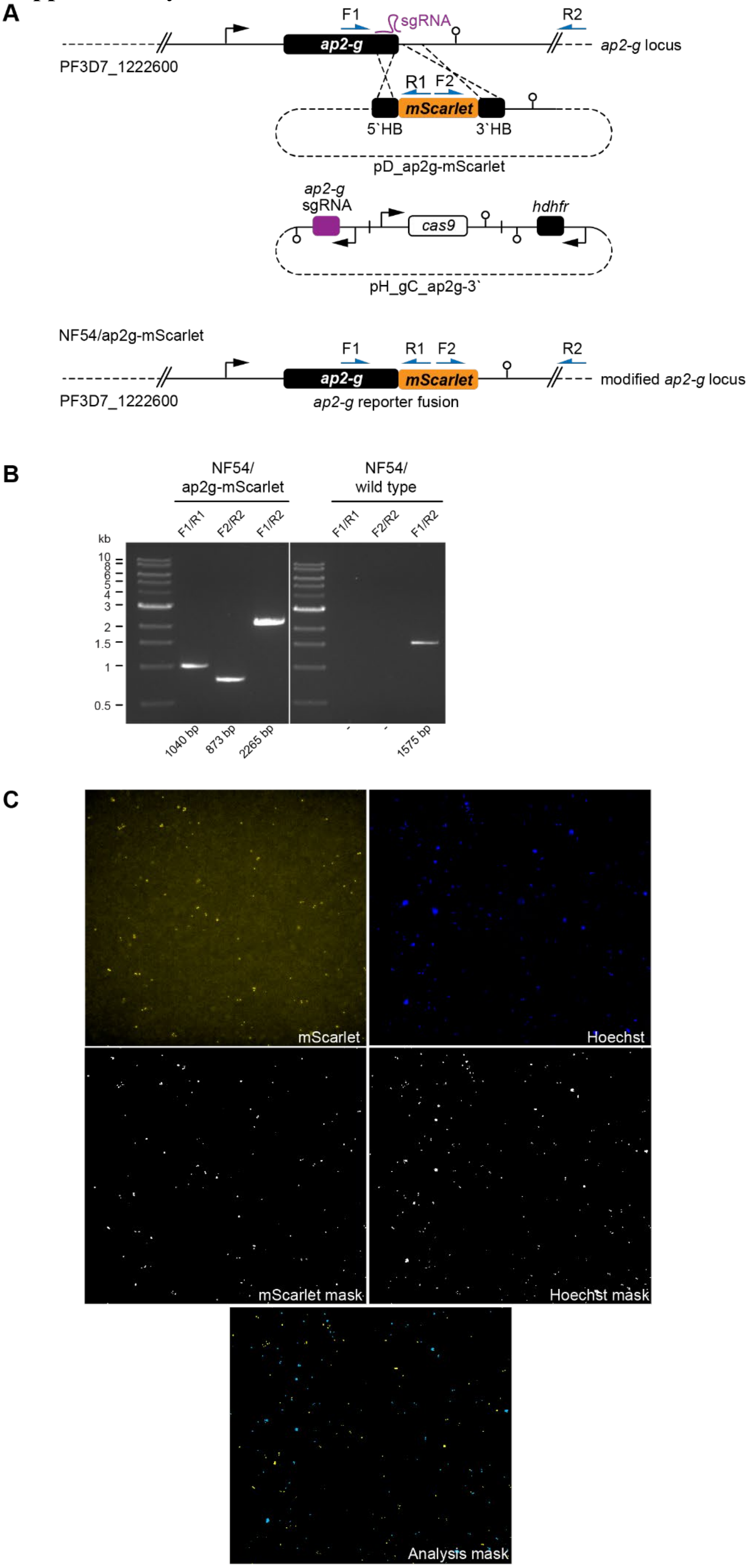

**Supplementary Figure 1. CRISPR/Cas9-based engineering of the NF54/ap2g-mScarlet parasite line.**

**(A)** Schematic maps of the wild type *ap2-g* locus (PF3D7_1222600), the CRISPR/Cas9 donor plasmid (pD_ap2g-mScarlet) and the suicide plasmid (pH_gC_ap2g-3') used to generate the modified *ap2-g* locus. Names and binding sites of the primers used for integration PCRs are indicated. **(B)** Integration PCRs performed on gDNA of NF54/ap2g-mScarlet parasites to confirm correct editing of the locus. PCRs performed on gDNA of NF54 wild type parasites served as control. Numbers at the bottom indicate the expected band sizes. **(C)** Example images from the HCI-based SCR quantification. Shown are raw images for AP2-G-mScarlet and Hoechst (top panel) and their respective masks produced by automated image analysis (middle panel). The bottom mask shows whether individual parasites were scored as sexually committed (yellow false-color) or asexually committed (blue false-color).

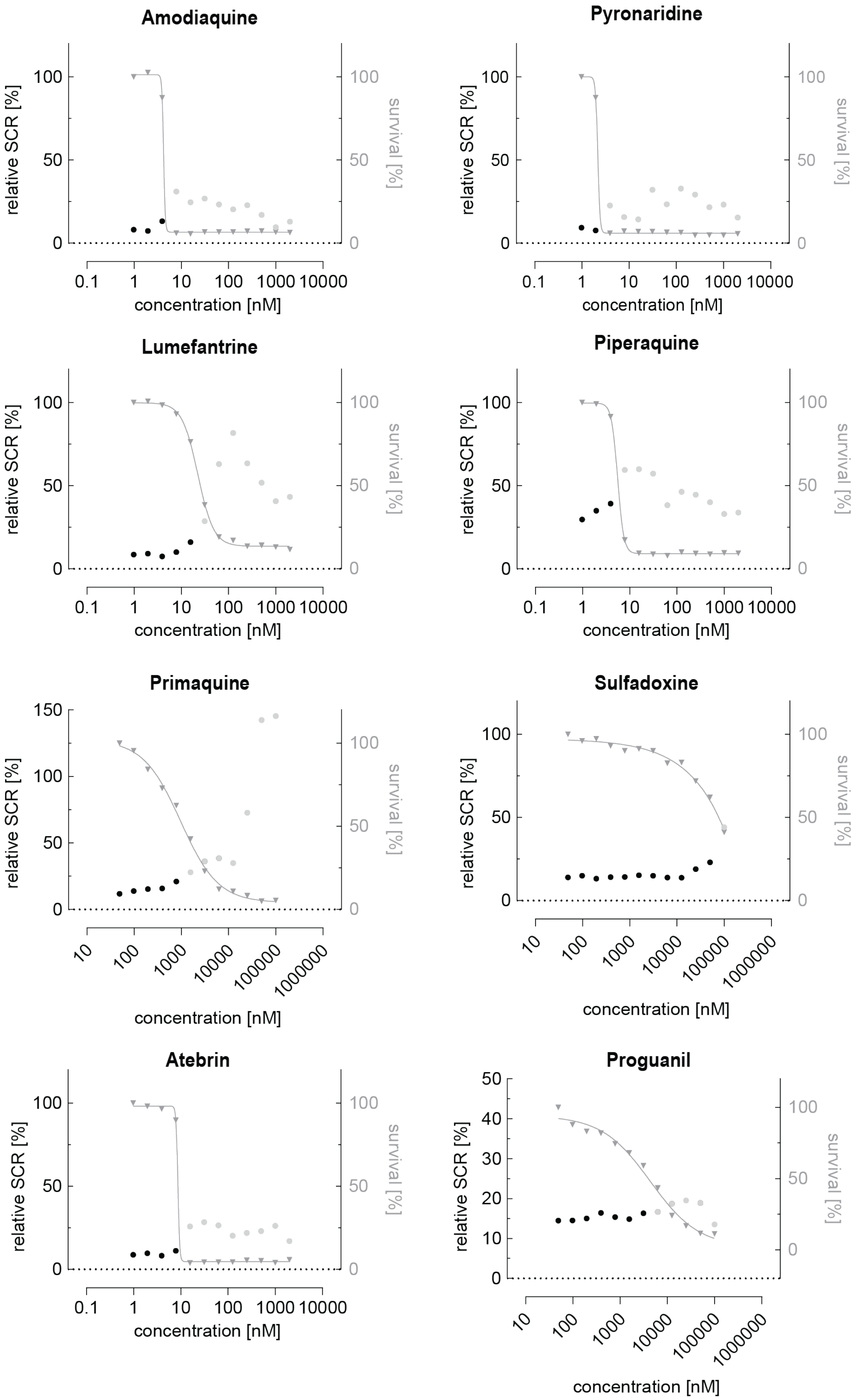


**Supplementary Figure 2. Dose-response relationship between antimalarial compounds and parasite sexual commitment.**

Antimalarials show a general trend towards induction of sexual commitment at growth-inhibiting concentrations. Mean parasite survival rates and SCRs are indicated by grey triangles and black bullets, respectively. Grey bullets represent SCRs at compound concentrations above the IC50. Values are normalized to the corresponding control conditions (-SerM for SCR) and (-SerM/choline for survival). n=1.

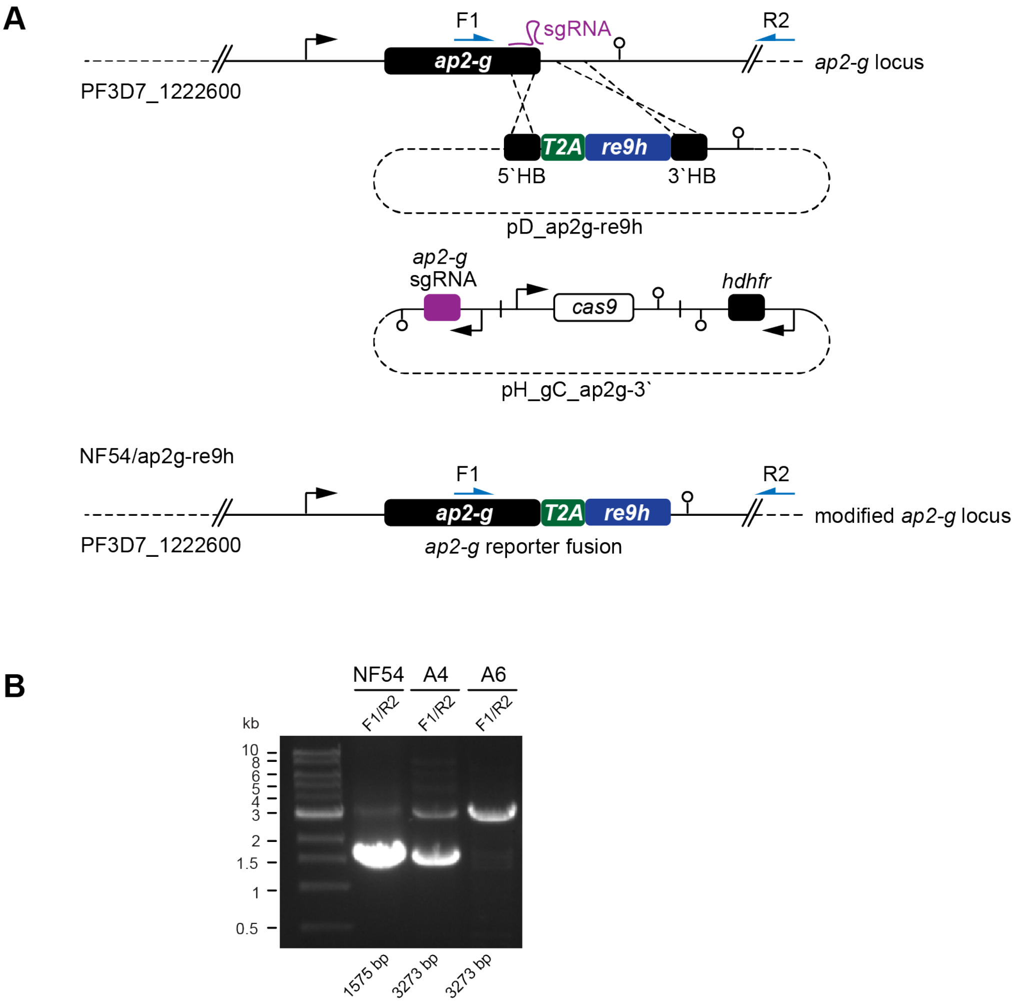


**Supplementary Figure 3. CRISPR/Cas9-based editing of the NF54/ap2g-re9h parasite line.**

**(A)** Schematic maps of the wild type *ap2-g* locus (PF3D7_1222600), the CRISPR/Cas9 donor plasmid (pD_ap2g-re9h) and the suicide plasmid (pH_gC_ap2g-3') used to generate the modified *ap2-g* locus. A sequence encoding the self-cleaving T2A peptide separates the *ap2-g* and *re9h* coding sequences. Names and binding sites of the primers used for integration PCRs are indicated. **(B)** Integration PCRs performed on gDNA of two NF54/ap2g-re9h parasite lines obtained from independent transfections (A4, A6) to confirm correct editing of the locus. PCRs performed on gDNA of NF54 wild type parasites served as control. Numbers at the bottom indicate the expected fragment sizes. The NF54/ap2g-re9h A6 line was used for all subsequent experiments.

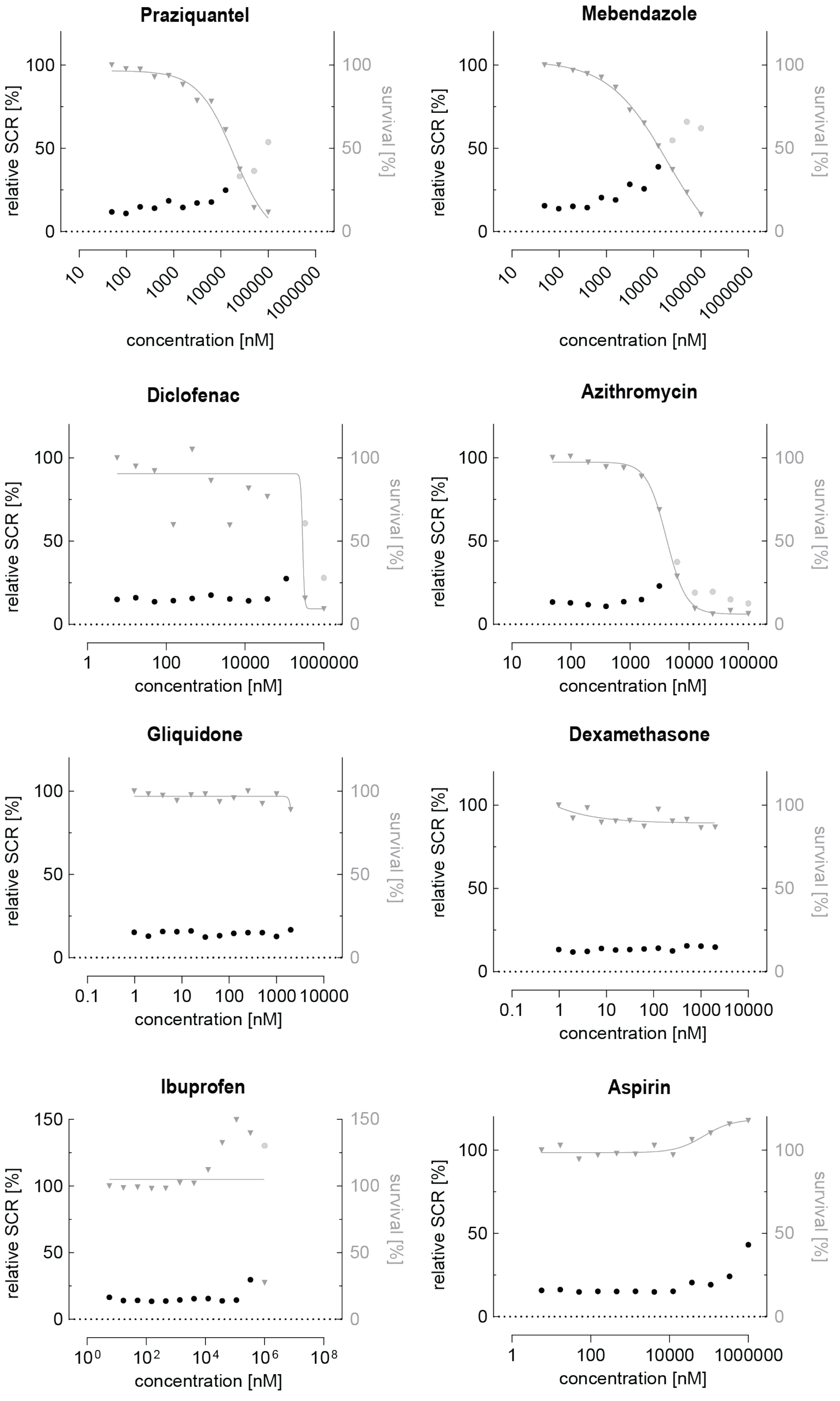


**Supplementary Figure 4. Dose-response relationship between commonly used drugs and parasite sexual commitment.**

The antihelminthics praziquantel and mebendazole show a general trend towards induction of sexual commitment at super-physiological concentrations that also affect parasite growth. The antipyretic diclofenac and the macrolide antibiotic azithromycin show a similar trend. The antidiabetic gliquidone and the corticosteroid dexamethasone do neither affect parasite growth nor SCR at the tested concentrations. Ibuprofen and Aspirin, two commonly used antipyretics, have a positive impact on parasite growth. Parasite survival rates and SCRs are indicated by grey triangles and black bullets, respectively. Values are normalized to the corresponding control conditions (-SerM for SCR) and (-SerM/choline for survival). n=1.

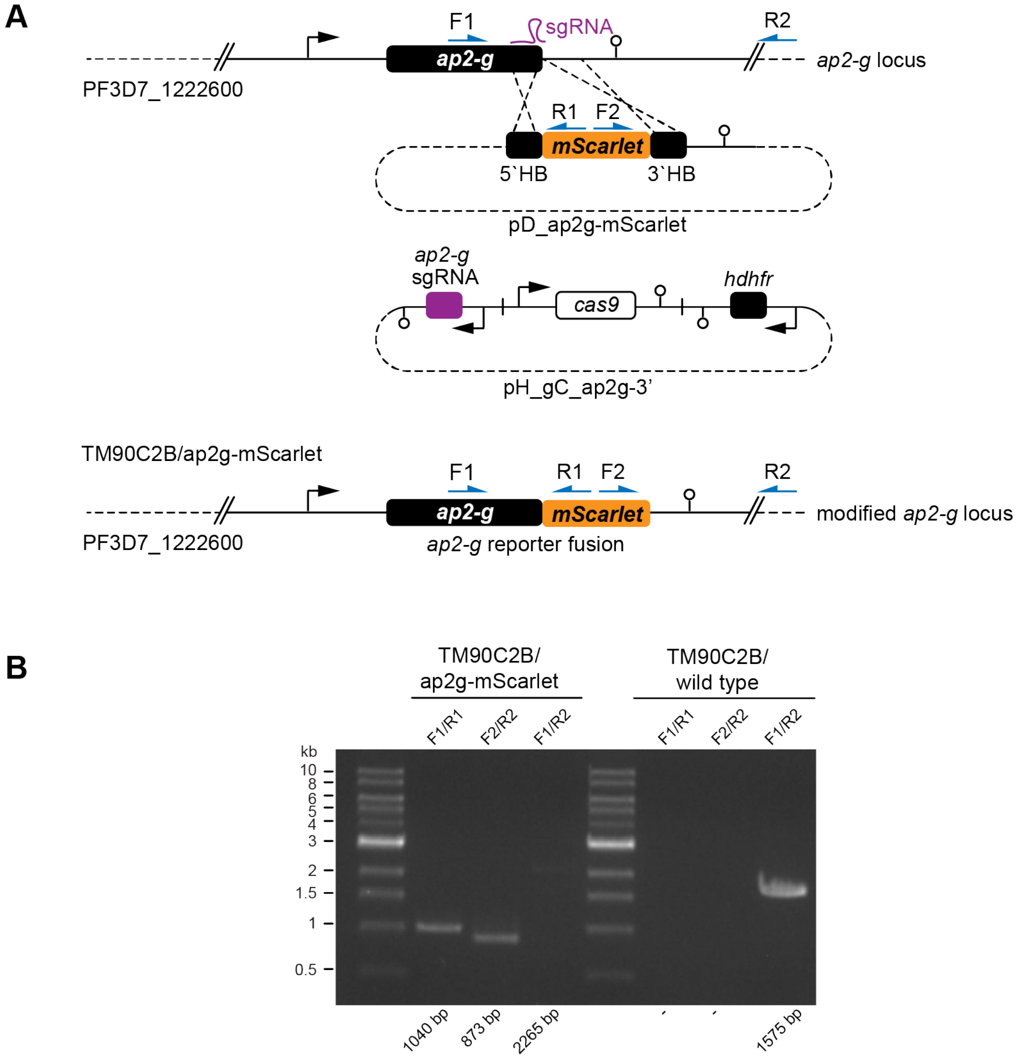


**Supplementary Figure 5. CRISPR/Cas9-based editing of the TM90C2B/ap2g-mScarlet parasite line.**

**(A)** Schematic maps of the wild type *ap2-g* locus (PF3D7_1222600), the CRISPR/Cas9 donor plasmid (pD_ap2g-mScarlet) and the suicide plasmid (pH_gC_ap2g-3') used to generate the modified *ap2-g* locus. Names and binding sites of the primers used for integration PCRs are indicated. **(B)** Integration PCRs performed on gDNA of TM90C2B/ap2g-mScarlet parasites to confirm correct editing of the locus. PCRs performed on gDNA of NF54 wild type parasites served as control. Numbers at the bottom indicate the expected fragment sizes.

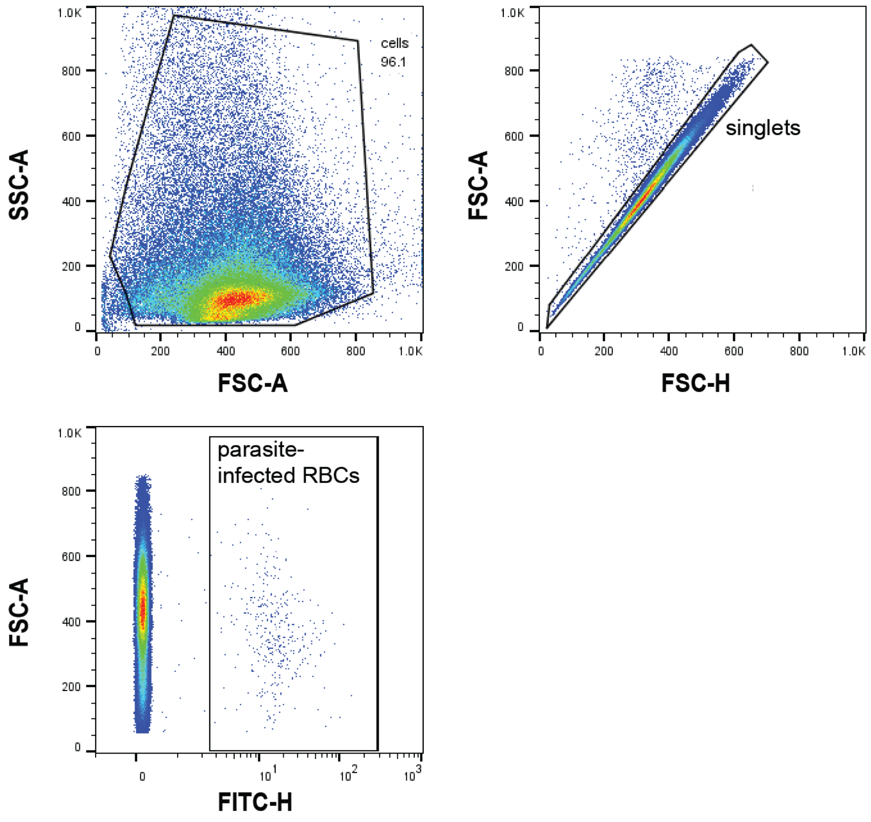


**Supplementary Figure 6. Flow cytometry gating strategy.**
Gating strategy of flow cytometry data. Representative flow cytometry plots of infected red blood cells. The first plot shows the gate to remove small debris (forward scatter vs. side scatter). The population 'cells' was then gated to remove doublets (forward scatter height vs. forward scatter area), keeping 'singlets' used for further gating. Finally, SYBR green fluorescence intensity was used to distinguish uninfected from infected RBCs (FITC vs forward scatter).

### Supplementary Tables

#### Table S1

Sequences of all oligonucleotides used in this study (5' to 3' orientation)

| **Cloning of pD_ap2g-mScarlet** | |
|---|---|
| mScarlet-F | gggaaacaacaggaatattggatccgcaggtagtaaaggtgaagcagttataaaag |
| mScarlet-R | gttatagggaatattcaaaatcatttatataattcatccattccacc |
| Ap2-g-3'-F | aatgattttgaatattccctataactaaatg |
| Ap2-g-3'-SfoI-R | gagaaaataccgcatcaggcctgtgtgttgagaaatttaaag |

| **Cloning of pD_ap2g-re9h** | |
|---|---|
| iso_re9h-F | gcaagatctgattttgaatattccctataactaaatgt |
| iso_re9h-R | tctacatctccacatgttaataaacttcctcttccttctccgctaccaatattcctgttgtttcccc |
| re9h-F | attaacatgtggagatgtagaagaaaatccaggaccaggtatggaggacgccaagaacatc |

| | |
|---|---|
| re9h-R | gaatattcaaaatcagatcttgccgcccttcttg |

| **PCR on gDNA** | |
|---|---|
| F1 | ggtaatgatggtgcattgag |
| R1 | gtaactgcacctccatcttc |
| F2 | gaacaatatgaaagaagtgaagg |
| R2 | gtaatacttattgtgtggaagg |

**Table S2**
ImageXpress modules used for automated image analysis (sequential steps).

| | | | |
|---|---|---|---|
| 1 | Setup | | |
| | | Image Names: | Channels: |
| | | DAPI | DAPI |
| | | TRITC | TRITC |

This step defines the channels used for analysis.

| | | | |
|---|---|---|---|
| 2 | Find round objects | | |
| | | Source | DAPI |
| | | Approximate Minimum With (µm) | 30 |
| | | Approximate Maximum With (µm) | 10000 |
| | | Intensity Above Local Background | 4000 |
| | | Result | Background DAPI |

This step identifies background signal in the DAPI channel.

| | | | |
|---|---|---|---|
| 3 | Find round objects | | |
| | | Source | TRITC |
| | | Approximate Minimum With (µm) | 4 |
| | | Approximate Maximum With (µm) | 50 |
| | | Intensity Above Local Background | 800 |
| | | Result | Background TRITC |

This step identifies background signal in the TRITC channel.

| | | | |
|---|---|---|---|
| 4 | Grow Objects | | |
| | | Source | Background TRITC |
| | | Grow by (pixels) | 10 |

| | | | |
|---|---|---|---|
| | | Result | Grow Objects TRITC |

This step expands the previously identified areas with background signal in the TRITC channel.

| | | | |
|---|---|---|---|
| 5 | Grow Objects | | |
| | | Source | Background DAPI |
| | | Grow by (pixels) | 50 |
| | | Result | Grow Objects DAPI |

This step expands the previously identified areas with background signal in the DAPI channel.

| | | | |
|---|---|---|---|
| 6 | Find Round Objects | | |
| | | Source | DAPI |
| | | Approximate Minimum With (µm) | 0.9 |
| | | Approximate Maximum With (µm) | 3.2 |
| | | Intensity Above Local Background | 4000 |
| | | Result | iRBCs |

This step identifies DAPI signal of infected red blood cells.

| | | | |
|---|---|---|---|
| 7 | Remove Marked Objects | | |
| | | Objects Source | iRBCs |
| | | Marker Source | Grow Objects DAPI |
| | | Result | iRBCs wo background |

This step removes areas with background signal from the DAPI channel.

| | | | |
|---|---|---|---|
| 8 | Remove Marked Objects | | |
| | | Objects Source | iRBCs wo background |
| | | Marker Source | Grow Objects TRITC |
| | | Result | iRBCs wo background DAPITRITC |

This step removes areas with background signal from the TRITC channel.

| | | | |
|---|---|---|---|
| 9 | Find Round Objects | | |
| | | Source | TRITC |
| | | Approximate Minimum With (µm) | 0.4 |
| | | Approximate Maximum With (µm) | 3.2 |
| | | Intensity Above Local Background | 800 |
| | | Result | Scarlet Signal |

This step identifies mScarlet signal from sexually committed parasites.

| | | | |
|---|---|---|---|
| 10 | Keep Marked Objects | | |
| | | Objects Source | iRBCs wo background DAPITRITC |
| | | Marker Source | Scarlet Signal |
| | | Result | Sexual parasites |

This step compares the background-corrected DAPI signal of infected red blood cells to the mScarlet signal of sexually committed parasites and only keeps double-positive parasites.

| | | | |
|---|---|---|---|
| 11 | Measure Mask | | |
| | Measurement Inputs: | Standard Area Value | 1 |
| | Objects to Measure: | Mask of Objects: | iRBCs wo background DAPITRITC |
| | | Image to Measure: | iRBCs wo background DAPITRITC |
| | Features within each Object: | Mask of Features: | Sexual parasites |
| | | Image to Measure: | iRBCs wo background DAPITRITC |

This step quantifies mScarlet-positive (sexually committed) and mScarlet-negative (asexual) parasites.
